# NDR2 Kinase Regulate Microglial Metabolic Adaptation and Inflammatory Response: Critical Role in Glucose-Dependent Functional Plasticity

**DOI:** 10.1101/2025.06.12.659313

**Authors:** Beatriz Fazendeiro, Ivo Machado, Anabela Rolo, Paulo Rodrigues-Santos, António Francisco Ambrósio, Paulo F. Santos, Hélène Léger

## Abstract

Diabetes and its retinal complication, diabetic retinopathy (DR), are a rapidly increasing health, societal and economic burden. Diabetic retinopathy is a complex disease with a chronic inflammatory component mediated by retinal microglial cells. Recent studies have demonstrated the importance of the Hippo pathway kinases, *Ndr1/Stk38* and *Ndr2/Stk38l*, in the regulation of macrophages, immune cells that share similarities with microglial cells. However, the role of NDR2 kinases in microglial inflammatory response and in the pathophysiology of diabetic retinopathy has not yet been uncovered. This study investigates the role of NDR2 kinase in microglial cells, particularly in response to high glucose (HG) conditions. Using CRISPR-Cas9, we downregulated *Ndr2* kinase in BV-2 microglial cells and analyzed the impact on cellular metabolism, phagocytosis and migratory capabilities.

We demonstrate that microglial cells expressed NDR2 kinase protein, especially in HG conditions, suggesting its importance in regulating microglial functions during hyperglycemia. *Ndr2* downregulated cells present a decreased basal respiration, indicating an impaired mitochondrial function. They also showed decreased metabolic flexibility to stress conditions, such as adaptation to HG conditions. Functionally, *Ndr2* downregulation led to decreased phagocytic capacity and migration of microglial cells, both cytoskeleton-based functions. Furthermore, *Ndr2* downregulation resulted in altered cytokine and chemokine secretion profiles. Notably, increased levels of pro-inflammatory cytokines such as IL-6, TNF, IL-17 and IL-12p70 were observed in *Ndr2* downregulated cells, even under normal glucose conditions. In conclusion, our findings indicate that NDR2 kinase is crucial for microglial metabolic adaptation to stress, such as high glucose exposure and for influencing microglial inflammatory responses. Therefore, NDR2 kinase plays a vital role in maintaining microglial functional plasticity in response to glucose variations, suggesting potential implications for neuroinflammatory processes in conditions like diabetic retinopathy.

**Graphical abstract:** 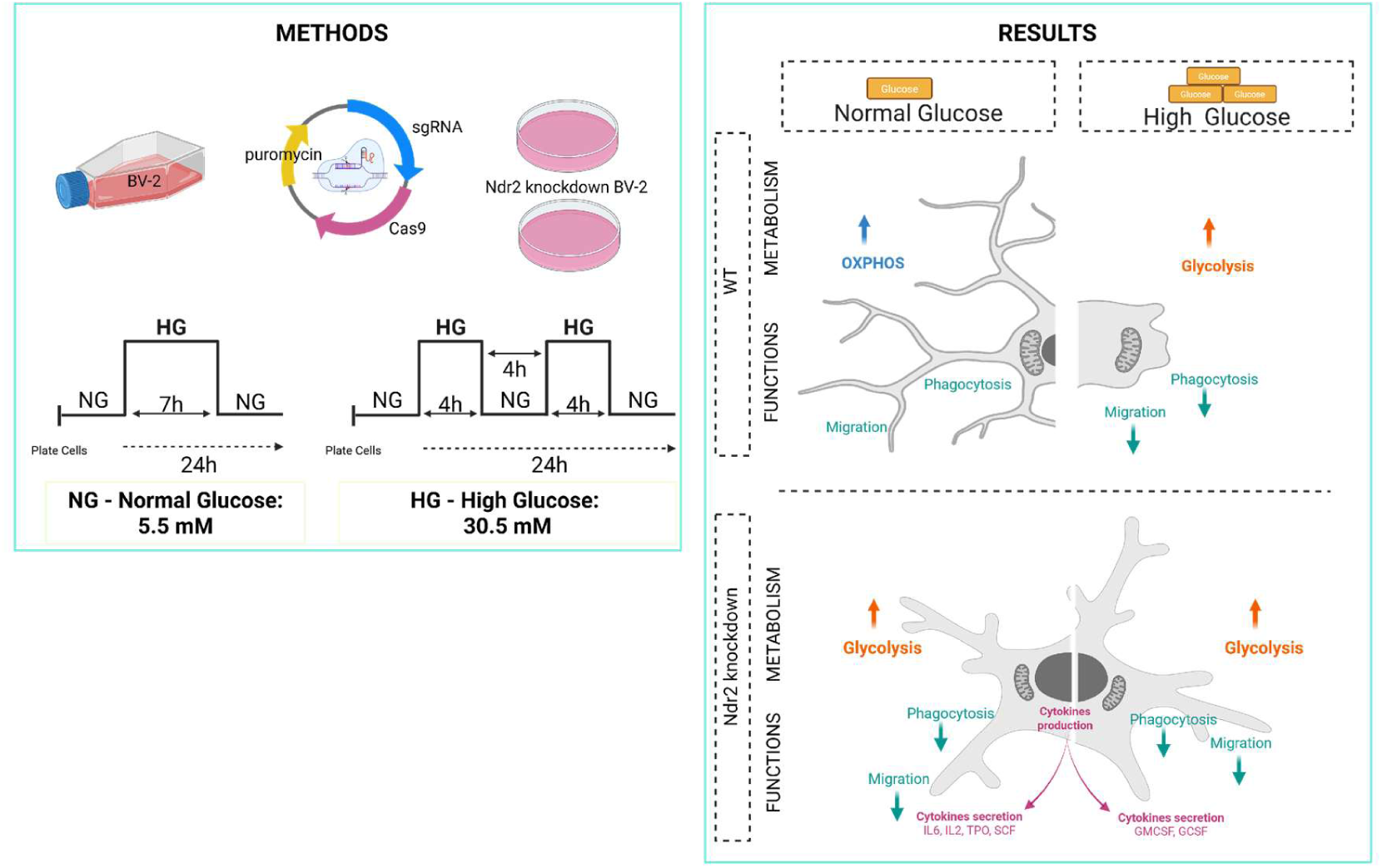

**Research in Context:** 

**What is already known about this subject?:**

- The pathogenesis of diabetic retinopathy (DR) involves chronic inflammation mediated by retinal microglial cells, which contribute to vascular damage and neurodegeneration.
- Microglial dysfunction under high glucose (HG) conditions exacerbates cytokine release and oxidative stress, driving DR progression.
- NDR kinases regulate inflammatory pathways in macrophages, but their role in microglia during DR was previously unexplored.

**What is the key question?:**

- How do NDR2 kinase regulate microglial inflammatory responses and functional adaptability in diabetic retinopathy?

**What are the new findings?:**

- NDR2 expression is upregulated in microglia exposed to HG.
- *Ndr2* downregulation in microglia impairs metabolic flexibility, phagocytosis, and migration.
- *Ndr2* downregulation disrupts cytoskeleton-dependent microglial functions, limiting their ability to adapt to metabolic stress.
- *Ndr2* downregulation in microglia increases pro-inflammatory cytokines (IL-17, TNF) and reduces anti-inflammatory factors (sTNFRI, VEGF), exacerbating inflammation.

**How might this impact clinical practice?:**

- Targeting Ndr2 signaling could emerge as a therapeutic strategy to modulate microglial-driven inflammation, potentially slowing DR progression and complementing existing glycemic control approaches.

## Introduction

Diabetes and its associated retinal disease - diabetic retinopathy (DR) - are a rapidly increasing health, societal and economic burden. About 93 million people worldwide have diabetic retinopathy (American Academy of Ophthalmology 2025) which affects 1 out of 3 persons with diabetes mellitus, making it the leading cause of vision loss among working-age adults.

DR is a complex disease with a chronic inflammatory component in the retina and many genetic and environmental factors have been considered to contribute to its development[1]. Impaired glucose metabolism, a key feature of diabetes, leads to biochemical changes that cause retinal vascular damage, a hallmark of the disease, also inducing the activation of the retinal microglial cells and the production of reactive oxygen species (ROS)[2]. Microglial cells are mononuclear phagocytes like macrophages, and the first line of defense of the nervous system. Activated microglia contribute to the progression of DR by releasing cytokines, chemokines, and other molecules that promote vascular leakage, abnormal blood vessel growth, and vascular and neural cell death in the retina[3, 4]. Understanding the role of microglial cells and inflammation in DR has led to research into potential therapies that target these cells and neuroinflammation. Strategies to modulate microglial activation or reduce inflammation could potentially slow or prevent the progression of DR, complementing traditional approaches focused on blood glucose control and vascular health.

The NDR (Nuclear Dbf2-Related) kinases (*Ndr1/Stk38* and *Ndr2/Stk38l*) are serine/threonine protein kinases that are evolutionarily conserved and play essential roles in regulating various cellular processes such as cell growth and apoptosis[5–7]. Dysregulation of NDR kinase signaling has been implicated in various diseases, including cancer, neurodegenerative disorders, and metabolic diseases[8, 9]. While the NDR kinases are not extensively studied in the context of inflammation compared to other kinases, there is emerging research linking NDR kinases to inflammation mediated by macrophages by regulating the pattern-recognition receptors and via their interactions with various signaling pathways such as the nuclear factor-κB[10, 11].

In the retina, we have previously established that *Ndr2* and *Ndr1* deletion in mice leads to a concomitant increase in apoptosis and cell proliferation of terminally differentiated neurons[12] similar to the phenotype observed in the canine early retinal degeneration (erd) disease[13–15]. Enrichment analyses of the differentially expressed genes (DEGs) from (*Ndr2*) erd dog retinas present an early activation of immune response genes, such as genes encoding constituents of NLRP3 inflammasome, and common components of IL1-β, IL-18 and TLR4 inflammatory pathways[16]. In mouse retinas, enrichment analyses of the DEGs point out that *Ndr2* deletion causes deregulation of retinal stress- and inflammation-related gene expression with enrichment of genes associated with oxidative stress, and cytoskeleton misregulation[12]. New causal analysis of the DEGs highlighted the putative role of the NDR2 kinase in the inflammation pathways mTOR, CXCR4 and eIF17. In neurons, mitochondrial dysfunction, oxidative phosphorylation, PI3K/AKT, mTOR, eIF2, eIF4 and p70S6K signaling lead to an inflammatory response and are known to be regulated by the microglia[17]. However, the exact functions and mechanisms of NDR kinases in retinal inflammation and in the context of diabetic retinopathy are not yet known.

This study aims to elucidate the role of NDR2 kinase as a new key player in microglial cells exposed to high glucose conditions. This will permit a better understanding of the regulation of microglial cells, and their surveillance and damage-sensing functions while exposed to metabolic stress, such as hypoglycemia in the pathology of diabetic retinopathy[18–20].

## Methods and Materials

### Animal care

All procedures involving animals were approved by the Animal Welfare Committee of the Faculty of Medicine of University of Coimbra (ORBEA 9-2022) and conducted in accordance with the European Community directive guidelines for the use of animals in laboratory (2010/63/EU) transposed to the Portuguese law in 2013 (Decreto-Lei 113/2013), and in agreement with the Association for Research in Vision and Ophthalmology statement for animal use.

### Mouse retinal microglial cell cultures

Primary retinal microglial cell cultures were obtained from 3 to 4-day-old C57BL/6J mice pups, as described previously[21, 22], with some modifications, as follows. The retinas were dissociated by trypsinization (0.25% trypsin supplemented with 0.05 mg/ml DNase I) followed by mechanical dissociation. The cell suspension was plated in uncoated T75 flasks (corresponding to eight retinas per flask) and maintained in Dulbecco’s Modified Eagle Medium/Nutrient Mixture F-12 (DMEM/F-12) with GlutaMAX™ (GIBCO) supplemented with 10% Fetal Bovine Serum (FBS, GIBCO), 100 U/ml penicillin, 100 μg/ml streptomycin (GIBCO) and 50 ng/ml of recombinant mouse CSF1 (Stemcell #78059.1). A mild trypsinization (0.25% trypsin) was performed every 3-4 days to remove active microglia cells and prevent cytokine overload of the culture. When the culture reached near confluency (∼2 weeks), cells were trypsinized and plated at a density of 0.5 × 10^4^ cells / well in µ-Slide 8 Well high (Ibidi #80801) coated with poly-D-lysine (0.1 mg/ml) and cultured at 37°C in a humidified atmosphere of 5% CO2. The purity of the culture (>90%) was assessed by immunocytochemistry with anti-Iba1 antibody.

### Human iPSC-derived microglial cell cultures

Macrophage/microglia progenitors were provided by Dr. Ana Luisa Cardoso (CNC, UC) and prepared as described by Haenseler et al[23]. Briefly, approximately 4 weeks after plating embryoid bodies in T175 flasks in the presence of X-VIVO15 medium (Lonza), supplemented with 100 ng/mL M-CSF (Invitrogen), 25 ng/ml IL-3 (R&D), 2 mM Glutamax (Invitrogen), 100 U/ml penicillin and 100 mg/ml streptomycin (Invitrogen), and 0.055 mM b-mercaptoethanol (Invitrogen), with fresh medium added weekly, macrophage/microglia progenitors start to emerge in the supernatant. These cells were collected by medium centrifugation and further differentiated into iPSC-derived microglial cultures. For that purpose, cells were counted and seeded at 3.0 × 10^4^ cells/well in fibronectin-coated Ibidi µ-Slide 8 Well plates, in DMEM/F12 medium supplemented with N2, 100 ng/ml IL-34 (R&D) and 10 ng/ml GM-CSF (R&D). The microglial progenitors were matured for 2 weeks, with medium changes 3 times per week, before being used for experiments.

### BV-2 cell Culture

BV-2 cells were cultured in Roswell Park Memorial Institute 1640 culture medium (RPMI; Gibco #11875093) supplemented with 10% (v/v) FBS, 1% (v/v) L-glutamine (ThermoFisher Scientific #25030024), and 1% (v/v) Penicillin-Streptomycin (Pen/Strep; ThermoFisher Scientific #15140122). Cells were then maintained in RPMI at 37 °C in a humidified atmosphere with 5% CO2 and kept until passage 27 (P27).

### High glucose assay

BV-2 cells were seeded at a density of 1.0 x 10^4^ cells / cm^2^ in Dulbecco’s Modified Eagle Medium low glucose (DMEM NG; Gibco 31600083) containing 5.5 mM glucose (1 g/l) and supplemented with 10% (v/v) FBS, 2mM L-glutamine, and 1% (v/v) Pen/Strep. This condition is called normal glucose condition (NG) or control condition (CT). After 16h-18h, in some of the wells, the culture medium was supplemented with 25 mM D-glucose (Sigma #G8270) in addition to the 5.5 mM already present in DMEM, reaching a final concentration of 30.5 mM (HG; 5 g/l) to simulate hyperglycemic conditions observed in diabetes.

The cells were exposed to HG for 7h (7h assay) or two times 4h with an interval of 4h between exposures in HG (12h assay). Then, BV-2 were kept in normal DMEM until the next morning for subsequent experiments (Figure 1).

**Figure 1.**
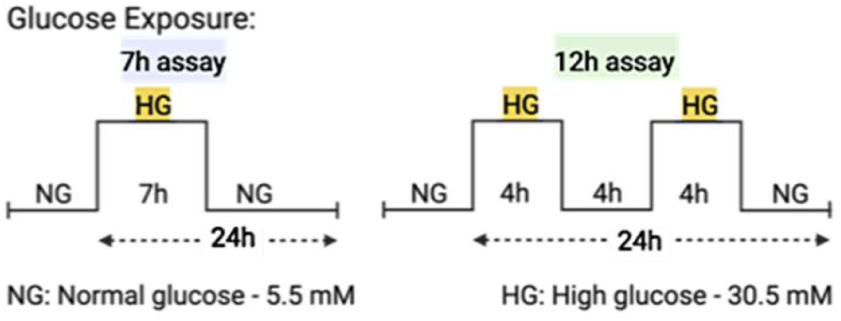
Schematic representation of the different experimental protocols used for exposure to high glucose levels. BV- 2 cell cultures were incubated in 5.5 mM glucose (normal glucose; NG or CT) or 30.5 mM glucose (high glucose; HG) for different periods: 7h (7h assay) or two times 4h HG with a break of 4h in between in NG (12h assay). (Created with BioRender)

### Plasmids, BV-2 transfection and clonal selection

#### Plasmid

*Ndr2/Stk38l* clonal downregulated BV-2 cell lines were generated by introduction of frameshift mutations in *Ndr2* exon 7 by CRISPR/Cas9 gene-targeting methods, as described[24]. The empty plasmid pX459, catalog number #62988, containing a puromycin resistance cassette and Cas9 nuclease, was obtained from Addgene. The all-in-one plasmid used for genome editing was obtained from Vector Builder (Spain) and contains one sgRNA targeting the *Ndr2* exon 7 (Supp. Table S1), designed using the CRISPR design tool CHOPCHOP (http://chopchop.cbu.uib.no/index. php)[25, 26] and inserted in the pRP[CRISPR]-Puro- hCas9-U6 vector allowing the co-expression of a puromycin resistance cassette and Cas9 nuclease. The vector ID is VB230329-1589djw, which can be used to retrieve detailed information about the vector on vectorbuilder.com.

The specificity of CRISPR/Cas9 editing depends on the sgRNA sequence and on the presence of a protospacer adjacent motif located next to the target sequence. The sgRNA sequence was chosen to target *Ndr2* exon 7 coding for the kinase domain while exhibiting the highest quality score by inverse likelihood of off-target binding based on the CRISPR design tool developed by Zhang Lab, MIT 2017 (http://crispr.mit.edu/)[24].

### Transfection

BV-2 cells were seeded into a T25 flask, at a density of 2.0 x 10^4^ cells / cm^2^ with 5 mL of RPMI medium and then maintained at 37°C in a humidified atmosphere with 5% CO2 for 18h. The following day, transfection with Lipofectamine 3000 (ThermoFisher Sc. #L3000001) was performed on cells (70-90% confluency) for 8h in Opti-MEM (Gibco #31985- 047), after which Opti-MEM was changed to RPMI.

### Selection and isolation of clonal cell lines

The morning of the next day, puromycin (3 µg/ml) was added to the transfected cells and kept for 30h to select the cells that had successfully incorporated the plasmid of interest. Once the cells reached a confluence of approximately 70%, the transfected cells were diluted to 100 cells in 12 ml of RPMI and seeded into a 96- well plate to obtain multiple clones that could be further expanded. 10 colonies (round and radiating from a central point) were selected and allowed to expand for 2–3 weeks. The puromycin concentration for the selection of the transfected BV-2 cells was established by performing resazurin assay after 24h, 30h and 48h exposure to 2, 3 and 5 µg/mL of puromycin. For this test, we used the pX459 plasmid, carrying a puromycin-resistance cassette (data not shown).

### *Ndr2/Stk38l* CRISPR BV-2 cells validation

DNA was isolated from BV-2 cells using ethanol precipitation. Sanger sequencing was used to identify insertion/deletion (Indels) caused by non-homologous end joining repair at the cut site in *Ndr2* exon 7 (see Supp. Table S1 for oligos). To validate *Ndr2* deletion and compare relative Ndr2 transcript levels in WT and the selected clones (clone 8, 13, 19, and 22), quantitative RT-PCRs (RT-qPCR) were conducted to probe mouse cDNA for transcripts that span *Ndr2* exons 13–14 (see Supp. Table S1 for oligos).

### Quantitative RT-qPCR

Total RNA was extracted using a column-based RNA purification kit (Zymo Quick-RNA MiniPrep Kit, R1055), following the manufacturer’s protocol for adherent cells. For cDNA synthesis, samples were considered suitable when the ratio values for both A260/A280 and A260/A230 were within the range of 1.8 to 2.2. cDNA was synthesized using the NZY First-Strand cDNA Synthesis Kit (Nzytech), following the manufacturer’s instructions. The resulting cDNA was diluted to a final concentration of 25 ng/µl. RT-qPCR experiments were done in compliance with standard MIQE (Minimum Information for Publication of Quantitative Real-Time PCR Experiments) guidelines[27]. The RT-qPCR reactions contained 50 ng cDNA, 1x iTaq Universal SYBR Green mix (Bio-Rad #r1725122) and 250 nM of each unlabeled forward and reverse primer. Reactions were performed in 96-well reaction plates using the StepOnePlus PCR System (Applied Biosystems). *Ywhaz* was found to be the most stable housekeeping gene and used for normalization and calculation using the ΔΔCT method[27]. Primers for RT-qPCR are listed in Supp. Table S1.

### SDS-PAGE and western blot

Cells from at least 5 independent cultures were washed three times with ice-cold phosphate-buffered saline (PBS) and then homogenized in ice-cold RIPA lysis buffer (50 mM Tris HCl, pH 7.4, 150 mM NaCl, 5 mM ethylenediamine tetra-acetic acid (EDTA), 1% Triton X-100, 0.5% sodium deoxycholate, 0.1% sodium dodecyl sulfate (SDS), 1 mM dithiothreitol supplemented with 10% protease inhibitors (Roche Complete Mini-EDTA free protease inhibitors, Roche # COEDTAF-RO).

Proteins (20 µg of total protein loaded) were separated on a 12% SDS-polyacrylamide gel and transferred onto a PVDF membrane. Membranes were blocked in 5% (w/v) low-fat dry milk / Tris-buffered saline with 0.1% (v/v) Tween 20 (TBS-T) for 1 h at room temperature (RT), treated with primary antibody in blocking buffer overnight (O/N) at 4°C and incubated for 1 h at RT in the secondary antibody. Chemiluminescence was detected using the western-bright ECL kit (Advansta #K-12043-D20). Band intensity was quantified through Image Lab Software (Biorad) and normalized to the intensity of loading protein control (calnexin). All commercially available antibodies used for immunoblots and immunofluorescence experiments are listed in Supp. Table S2.

### Alamar blue assay

Twenty-four hours after starting the HG assays, cell viability was assessed using resazurin dye (7-hydroxy-3H-phenoxazin-3-one 10-oxide; stock at 500 µM), diluted in PBS at a final concentration of 50 µM. Following a 4h incubation at 37°C and 5% CO2, the absorbance at 570 nm and 620 nm was measured by a multimode microplate reader (Biotek Synergy HT) to assess the conversion of resazurin to fluorescent resorufin. By analyzing the relative absorbance values, proportional to the amount of metabolically active live cells present in the culture, the percentage of cell viability and the metabolic status of BV-2 cultures can be determined, as follows:

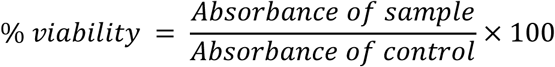

### Immunocytochemistry

Cells were washed with PBS and fixed with 100% methanol at -20°C for 10 min at RT. Subsequently, the cells were permeabilized and blocked with 3% bovine serum albumin (BSA) and 0.25% Triton X-100 in PBS for 30 min, treated with the primary antibody in blocking buffer O/N at 4°C and incubated for 1h at RT in secondary antibody and 0.5 µg/ml DAPI (Invitrogen #D1306).

### Microscopy and imaging/quantification

Immunofluorescence microscopy as conducted using either a widefield imaging microscope (Zeiss Axio Observer.Z1 inverted microscope equipped with a Colibri 7 LED light source, CCD digital camera (AxioCam HRm) and Zen Blue 2012 software) with a 20× (Plan-Apochromat, 0.8 NA) or an 40× (LD Plan-Neofluar, 0.6 NA Corr) objective, or a laser scanning confocal microscope (LSM 710 Axio Observer.Z1 microscope with QUASAR detection unit and Zen Black 2010 software) with a 63× (Plan-Apochromat, 1.4 NA oil) objective. All the equipment was from Carl Zeiss. All experimental conditions within a particular experiment were processed simultaneously, and imaging settings (exposure time and laser power) were conserved. An average of 5 images per coverslip were selected to quantify Ndr2, and analyzed using ImageJ, normalized to the number of cells per image, and the mean fluorescence intensity from these five images was used to determine the result for each condition.

CTCF = Integrated Density − (Area of selected cells × Mean fluorescence background)

### Phagocytic activity assay

Fluorescent yellow-green latex beads (Sigma #L1030-0001) were added to the cell medium to reach a final concentration of 0.0025% and incubated for 1h at 37°C with 5% CO2. The cells were washed three times with PBS to remove the non-phagocytized beads and fixed with 100% methanol at -20°C for 10 min. BV-2 cells were stained with TRITC-conjugated phalloidin (1:500; Sigma-Aldrich #P1951) and nuclei were stained with 0.5 µg/ml DAPI (Invitrogen #D1306). Cells were observed with a Carl Zeiss widefield microscope (Zeiss Axio Observer.Z1 inverted microscope), and five random fields were acquired from each condition. The analysis was performed through ImageJ. The phagocytic efficiency was calculated based on the following formula, with x being the number of beads[28]

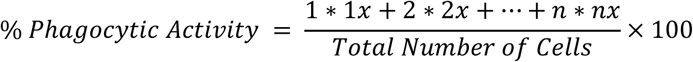

### Boyden chamber assay

24h prior to the assays in which BV-2 cells were exposed to glucose, the medium of BV-2 cells was replaced with DMEM containing 0% FBS. The next day, BV-2 cells were seeded into Transwell cell culture inserts (8 µm pore diameter, Millipore) at a density of 1 × 10⁴ cells/cm² in either NG or HG conditions. At the end of the experiments, cells were fixed in 4% PFA with 4% sucrose, and after 15 min, cells on the upper side of the transwell were removed with a cotton swab. Nuclei were stained with DAPI (1:2000) for 20 min to allow cell counting. The membranes were carefully removed from the inserts with a blade and mounted on glass slides with Dako Fluorescent Mounting Medium (Agilent #S302380-2). The samples were observed in an inverted fluorescence microscope (Zeiss Axio Observer.Z1 inverted microscope) and five fields were acquired from each membrane. The number of migrated cells was counted.

### Proliferation assay (EdU)

Cell proliferation was evaluated using the EdU staining proliferation kit (Abcam #ab222421), following the manufacturer’s protocol. EdU was added to the cells at a concentration of 10 µM Five random fields were acquired from each coverslip, the number of Edu-positive cells was counted, and the results were normalized to the total number of microglia. The results are presented as the percentage of the WT control.

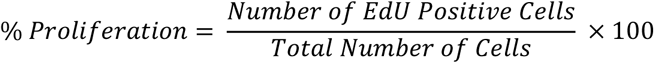

### Flow cytometry

#### Cell death assay

Cell death was assessed using flow cytometry through the double staining with annexin V (AV) and 7-aminoactinomycin D (7-AAD, Biolegend). After HG exposure, the cells were washed with PBS by centrifugation at 400× g for 5 min. After that, the cells were stained with AV and 7-AAD and analyzed as described[29]. Briefly, 1.0 × 10^6^ cells were collected and washed with PBS, centrifuged at 500× g for 5 min, resuspended in 100 μl of AV binding buffer and incubated with 2.5 μl of AV and 5 μl of 7-AAD for 15 min in the dark at RT. Then, cells were diluted in 300 µl of AV binding buffer and analyzed in a FACSCalibur flow cytometer (Becton Dickinson). At least 25000 events were acquired using CellQuest software (Becton Dickinson) and analyzed using Kaluza Analysis Software (Beckman Coulter). Results represent the mean ± SEM of percentage of each cell population for 3-4 independents experiments: viable cells (AV−/7-AAD−), initial apoptotic (AV+/7-AAD−), late apoptotic/necrotic (AV+/7-AAD+), and necrotic cells (AV−/7-AAD+).

### Cytokines

Cells (5 x 10^5^) were seeded into a 10 cm dish and exposed to HG or lipopolysaccharide (LPS, 100 ng/mL) for 24 h. After 20h, 5 µl of Brefeldin A (Sigma) was added to the culture to inhibit the secretion of cytokines and allow better detection by flow cytometry. Following this, cells were gently detached, centrifuged for 5 min at 230× g, and resuspended in PBS for flow cytometry analysis.

For surface marker detection, cell pellets were incubated with fluorescent conjugated antibodies against CD11b-V500 (BD Pharmingen, clone M1/70, RRID:AB_398535), P2γ12-APC (Biolegend, clone S16607D, RRID:AB_2721468), TNFa-PE (Biolegend, clone TN3-19.12, RRID:AB_315418), and IL-17a-V450 (BD Pharmingen, clone N49-653, RRID:AB_1727539). The control conditions include unstained cells, and single labeling controls compensation of the flow cytometer. The work conditions were categorized into two tube groups: i) extracellular labeling without permeabilization (CD11b-V500, P2γ12-APC) and ii) combined extracellular and intracellular labeling with permeabilization. The antibodies for extracellular labeling were added in all tubes, thoroughly mixed, and then incubated for 15 min in the dark at RT. After incubation, one PBS wash was performed, and cells of the first group were resuspended in PBS for flow cytometry reading. For tubes of the second category, 100 µl of Fix and Perm A solution (Invitrogen) was added. After 10 min incubation, one PBS wash was performed followed by 20 min incubation in 100 µl of the intracellular antibodies (TNFa-PE, IL-17a-V450) diluted in Fix and Perm B solution (Invitrogen), in the dark and at RT. After incubation, cells were washed twice with PBS buffer and acquired in an 8-color flow cytometer, BD FACSCanto II (BD Biosciences). Data analysis was performed using FlowJo v.10.7 (BD Biosciences).

### Seahorse analysis

Cells were seeded in an Agilent Seahorse 24-well XF culture microplate at a density of 0.6 x 10^4^ cells/well in 250 µl of DMEM NG, allowed to adhere for 24h in a humidified incubator with 5% CO2 at 37°C. Cells were subsequently treated with 500 µl of DMEM HG or DMEM NG for 7h, and then cultured in 500 µl of DMEM NG until the following day. The day prior to the Seahorse experiment, the Seahorse XFSensor Cartridge was hydrated and calibrated with 1 ml of Seahorse XF Calibrant Solution in a non-CO2 37°C humidified incubator.

Mitochondrial functionality and ATP production of BV-2 cells were analyzed using 1.5 µM oligomycin (Sigma-Aldrich #O4876), 5 µM of BAM15 (Sigma-Aldrich #SML1760), 0.5 µM of rotenone (Sigma-Aldrich #R8875) and 1 µM of antimycin A (Sigma-Aldrich #A8674). Each experiment was performed for at least 3 independent biological replicates. Data was normalized based on protein content using the sulforhodamine B (SRB) assay[30].

### Mouse cytokine antibody array

Mouse cytokine antibody arrays (Membrane, 22 Targets, Abcam #ab133993) were used to profile the following cytokines secreted in the medium of WT and Ndr2 downregulated BV-2 cells, in NG and HG conditions: GCSF, GM-CSF, IL-2, IL-3, IL-4, IL-5, IL-6, IL-9, IL-10, IL-12 p40/p70, IL-12p70, IL-13, IL-17, IFN-gamma, MCP-1, MCP-5, RANTES, SCF, sTNF RI, TNF, Thrombopoietin or TPO, VEGF. Each sample is the pool of n=3-6 supernatants, vol/vol. The experiment was carried out according to the manufacturer’s instructions. Briefly, antibodies-coated membranes were first incubated for 30 min with 2 ml of blocking buffer. After 30 min, the blocking buffer was replaced with 1 ml supernatant from WT 7h CT, WT 7h HG, Clone 19 7h CT and Clone 19 7h HG BV-2 cells samples. Membranes were incubated overnight at 4°C with mild shaking. The next day, the membranes went through a large volume wash, 3 washes with the Wash Buffer I and 2 washes with the Wash Buffer II before being incubated with 1 ml biotin-conjugated antibodies overnight at 4°C with mild shaking. Lastly, the membranes were washed as before and incubated with HRP-conjugated streptavidin (overnight at 4°C with mild shaking), then revealed (< 10 min) using a chemiluminescence substrate monitored on a VersaDoc Imager/Quantity One software (Bio-Rad).

## Results

### NDR kinases are expressed by microglial cells

To demonstrate the expression and localization of native NDR kinases in microglial cells, we performed immunocytochemistry using commercially available antibodies targeting respectively the N-terminus (aa 1-100; NDR1/2 antibody (E-2) #sc-271703) and the C-terminus (aa 380-460; NDR2 antibody #STJ94368) of the human NDR2 kinase on human iPSC-derived microglial cultures (iMGs), BV-2 immortalized microglial cells and mouse primary retinal microglial cultures[12]. In both cultures the microglia phenotype was validated by immunocytochemistry using antibodies against markers of several cell types: i) neuronal marker NeuN (negative staining, data not shown), ii) Müller cell marker vimentin (negative staining, data not shown), iii) astrocytes marker GFAP (Fig 2A-B), and iv) microglial cell marker IBA1 (Fig. 2A-B).

**Figure 2.**
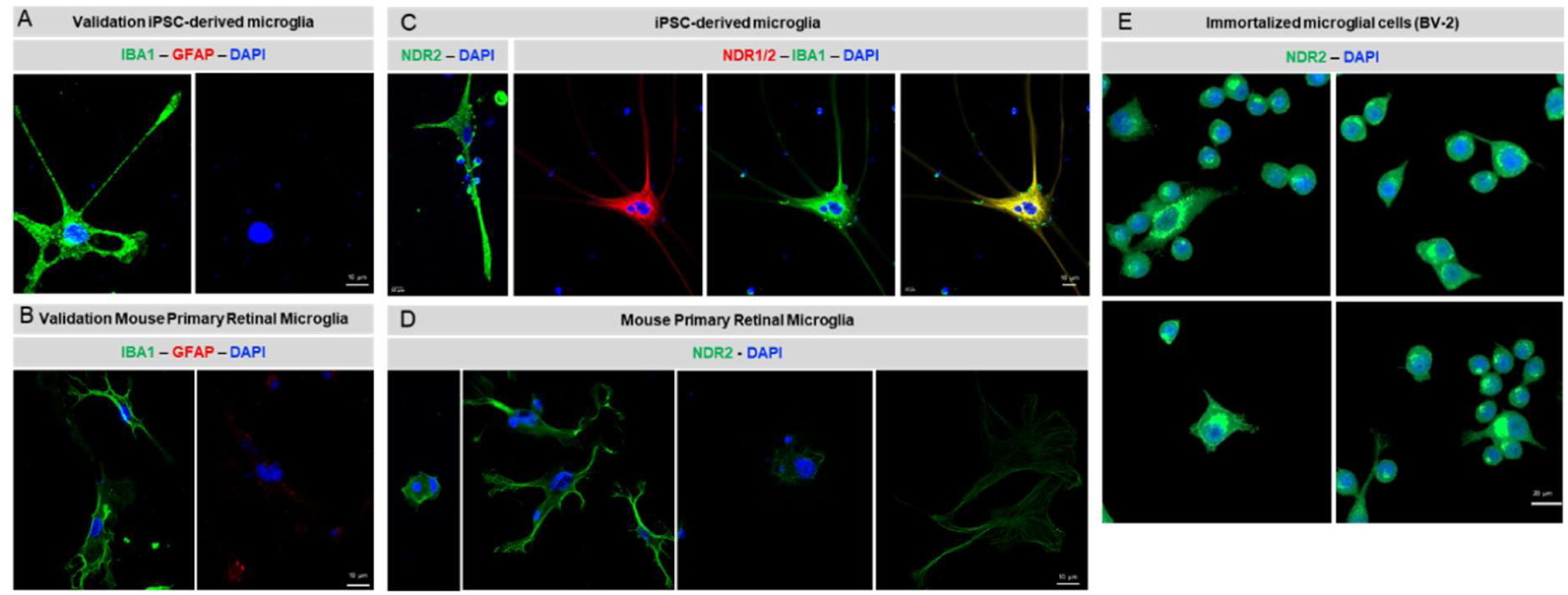
NDR2 expression in human and mouse microglial cells. (A, B) The microglia phenotype was validated in both human iPSC-derived microglial cultures and mouse primary microglial cultures by immunocytochemistry. IBA1: microglial cell marker (green); GFAP: astrocyte and reactive Müller cell marker (red). (C – E) Immunocytochemistry against NDR2 (green) in human iPSC-derived microglial cultures, mouse primary microglial cultures and BV-2 cell cultures. DAPI: nuclei staining (blue).

We were able to stain both NDR kinases in human microglial cells (Fig. 2C), while only the NDR2 antibody gave positive staining in mouse primary (Fig. 2D) and immortalized microglial cells (Fig. 2E). As shown in Figure 2C, NDR1 and NDR2 colocalized with the cytoskeletal protein IBA1 (ionized calcium binding adapter protein 1) in human iPSCs-derived microglial cells. In contrast, NDR2 staining in mouse primary retinal microglia was primarily localized mainly at the cell periphery and at the tips of microglial processes (Fig. 2D). NDR2 is localized in the cytoplasm in a peri-nuclear fashion in the immortalized microglial cell line BV-2. IBA1 is an actin-crosslinking protein, expressed specifically in microglial cells and macrophages, and it plays a crucial role in microglial cell migration, membrane ruffling, phagocytosis, and remodeling during immunological surveillance. In summary, the data shown in Figure 2 indicate that both NDR kinases are expressed by microglial cells and that NDR2 is expressed in microglial cells in the cytoplasm in a peri-nuclear fashion, at the cell periphery and at the tips of microglial processes.

### NDR2 protein expression is upregulated in microglial cells exposed to HG

To determine the effect of HG conditions on NDR2 protein levels, BV-2 cells were exposed to 30.5 mM of glucose for 7h or two times 4h with a 4h period in between exposed to NG (named the 12h assay), and western blot was performed. The results were normalized for calnexin, a loading control protein[31]. The exposure of BV-2 cells to HG conditions in the 7h assay (CT: 24.0 ± 4.4 a.u.; HG: 83.0 ± 19.1 a.u.) and in the 12h assay (CT: 26.1 ± 6.9 a.u.; HG: 64.2 ± 10.1 a.u.) caused a significant increase in NDR2 protein expression (Fig. 3A).

**Figure 3.**
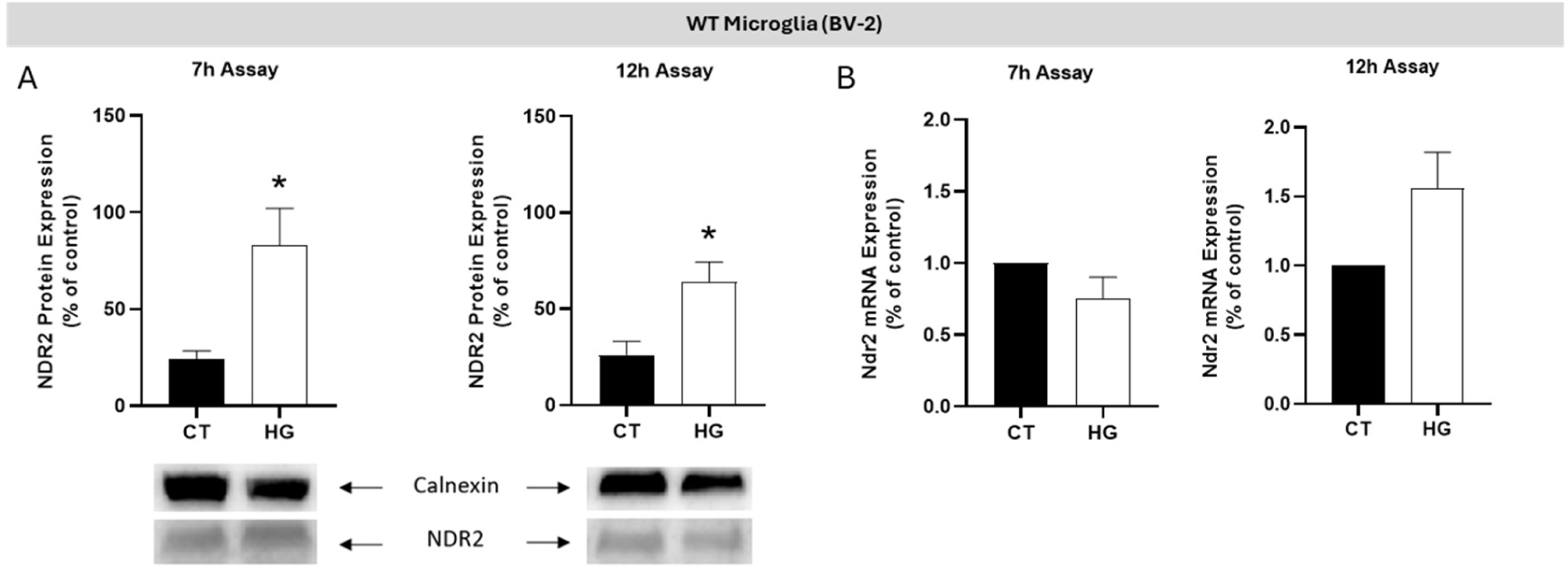
High glucose upregulates the NDR2 protein expression in BV-2 microglial cells. (A) The protein levels of NDR2 in BV-2 cells were exposed to 5.5 mM glucose (normal glucose condition; CT) or 30.5 mM glucose (HG) in the 7h assay and the 12h assay were evaluated by western blot analysis. Representative Western Blots are presented below the graphs. The results are expressed in arbitrary units (a.u.) ± SEM, and statistical analysis was assessed with a student’s t-test after confirmation of a Gaussian distribution; * p ≤ 0.05: HG compared with CT; n=6-7. (B) The mRNA levels of *Ndr2* in BV-2 cells exposed to 5.5 mM glucose (CT) or 30.5 mM glucose (HG) in the 7h assay and the 12h assay were evaluated by qRT-PCR analysis. The results are expressed as percentage of control ± SEM, and statistical analysis was assessed with a student’s t- test; n=5.

To determine the impact of HG exposure on *Ndr2*mRNA levels in BV-2 cells, we conducted a qRT-PCR analysis. As shown in Figure 3B, no alterations were observed in the expression of Ndr2 mRNA for the 7h HG exposure (HG: 80.0 ± 0.1 % of CT) when compared to CT conditions. However, a tendency towards an increase was found when cells were exposed to the 12h assay (HG: 160.3 ± 34.0 % of CT, p= 0.097). We also evaluated the impact of HG exposure on Ndr1 mRNA levels in BV-2 cells by qRT-PCR analysis and we found no alterations in the expression of Ndr1 for the 7h HG (HG: 80.8 ± 23.1 % of CT) and 12h HG exposure (HG: 120.3 ± 51.0 %) when compared to CT conditions (Supp. Fig. S1). In summary, these results suggest that HG conditions upregulate the NDR2 protein level but not NDR1 protein level in microglial cells.

To specifically investigate the role of NDR2 kinase in microglial cell responses in the context of HG exposure, we conducted a CRISPR-Cas9 lipofectamine transfection of the all-in-one plasmid containing a sgRNA against the exon 7 of the Ndr2 gene to induce a knockout of *Ndr2* in BV-2 cells.

### CRISPR-Cas 9 induced a downregulation of the Ndr2 gene in BV-2 cells

We transfected early passages BV-2 cells (p7) with the all-in-one plasmid VB230329-1589djw, targeting *Ndr2* exon 7 for 24 h. followed by puromycin selection for 30 h. After puromycin selection, we set up approximately 40 clonal cultures. Based on the phenotype of the clonal populations, we selected 10 of them for genomic analysis to check for insertion/deletion (Indels) after non-homologous end-joining (NHEJ) repair events using Sanger sequencing. We were able to classify the clones into three categories, each presenting an indel in the region near the PAM sequence (CCA in blue, Table S3). We chose to work with one clone of each of the distinct categories that were most likely to generate possible KOs: clone 13, 19 and clone 22. Two validation assays were performed to confirm that, indeed, we had a disruption of the *Ndr2* gene. Firstly, we assessed the *Ndr2* mRNA levels by qRT-PCR in the KO cells to compare with WT BV-2 cells. As we can see in Figure 4A, all clones presented a decrease in the mRNA levels when compared to the control group (Clone 13: 50 ± 10 % of CT; Clone 19: 23.5 ± 4.6 % of CT; Clone 22: 26.5 ± 2.9 % of CT) (see Supp. Table S3 for BV-2 *Ndr2* Clones). Clones 19 and 22 showed a greater reduction in *Ndr2* mRNA levels compared to clone 13, which led to their selection for further analysis. We performed immunocytochemistry on clones 19 and 22 to evaluate the NDR2 protein expression under CT condition using the commercially available antibody targeting the C-terminus (aa 380-460; NDR2 antibody #STJ94368) of the human NDR2 kinase (Fig. 4B). We also observed a decrease in the NDR2 protein levels in both selected clones when compared to WT BV-2 expression (Clone 19: 5.2 % of WT; Clone 22: 40.9 % of WT). Since we have amplification signals considering both qRT-PCR results and NDR2 expression from immunocytochemistry, we cannot consider that these clones present a real KO of the *Ndr2* gene. However, there is a clear downregulation of the Ndr2 gene for both clones.

**Figure 4.**
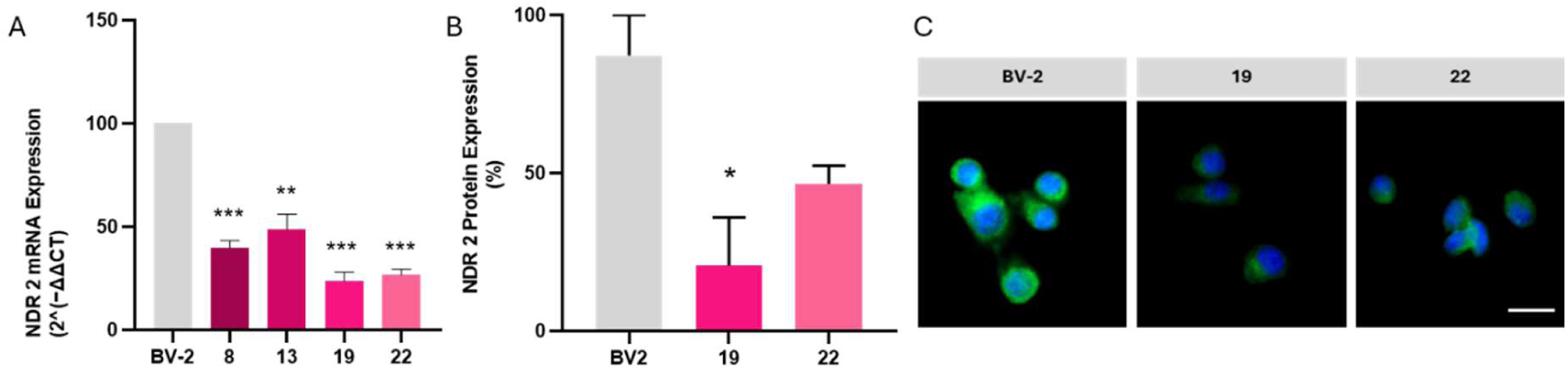
Ndr2 downregulated BV-2 clones. (A) The mRNA levels of *Ndr2* were evaluated by qRT-PCR in BV-2 (WT) and *Ndr2* downregulated BV-2 (Clones 8, 13, 19, and 22) cultured cells. The results are expressed as percentage of control ± SEM (n=2) and statistical analysis was assessed with one-way ANOVA test; * p ≤ 0.05 compared with WT CT, * p ≤ 0.05, compared with WT CT; ** p ≤ 0.005, compared with WT CT, *** p ≤ 0.001, compared with WT CT. (B) The expression of NDR2 protein was evaluated by immunocytochemistry analysis in BV-2 (WT) and *Ndr2* downregulated BV-2 (Clone 19 and Clone 22) cultured cells. These results are expressed as percentage of control of the means of fluorescence intensity (MFI) and statistical analysis was assessed with a one-way ANOVA test; * p ≤ 0.05, compared with WT CT; (n=2; 40 cells per independent experiment).

### High glucose and NDR2 downregulation effects on cell viability and metabolism

To assess the effects of *Ndr2* downregulation on the viability of BV-2 cells, we conducted an Alamar blue assay, which relies on resazurin reduction to resorufin. A non-statistically significant decrease in resazurin reduction was observed in both *Ndr2* downregulated BV-2 cells when compared to WT BV-2 cells (Clone 19 7h CT: 77.6 ± 12.0% control; Clone 22 7h CT: 68.9 ± 6.5% of control) (Fig. 5A). When WT cells were exposed to HG, no statistically significant alterations were registered compared to the CT condition (WT 7h HG: 83.8 ± 5.8% of WT CT). However, when *Ndr2* downregulated cells were exposed to 7h incubation with HG, a significant decrease was detected when compared to WT cells in CT conditions (Clone 19 7h HG: 67.9 ± 9.1% of WT CT; Clone 22 7h HG 67.0 ± 5.8% of WT CT). In the case of the 12h assay, only clone 19 exhibited a significant decrease in resazurin reduction when exposed to HG: WT 12h HG: 95.3 ± 4.1% of WT CT; Clone 19 12h CT: 57.6 ± 9.7% of WT CT, Clone 19 12h HG: 48.8 ± 12.8% of WT CT; Clone 22 12h CT: 77.1 ± 14.8% of WT CT; Clone 22 12h HG 75.9 ± 18.5% of WT CT) (Fig. 5B). Additionally, we incubated cells with 25 mM of mannitol (M) along with 5.5 mM of D-glucose already present in cell culture medium, being mannitol used as an osmotic control to ensure that the observed effects from HG exposure were not due to osmotic fluctuations. Our results demonstrate that mannitol did not affect the viability of the cells (Supp. Fig. S2).

**Figure 5.**
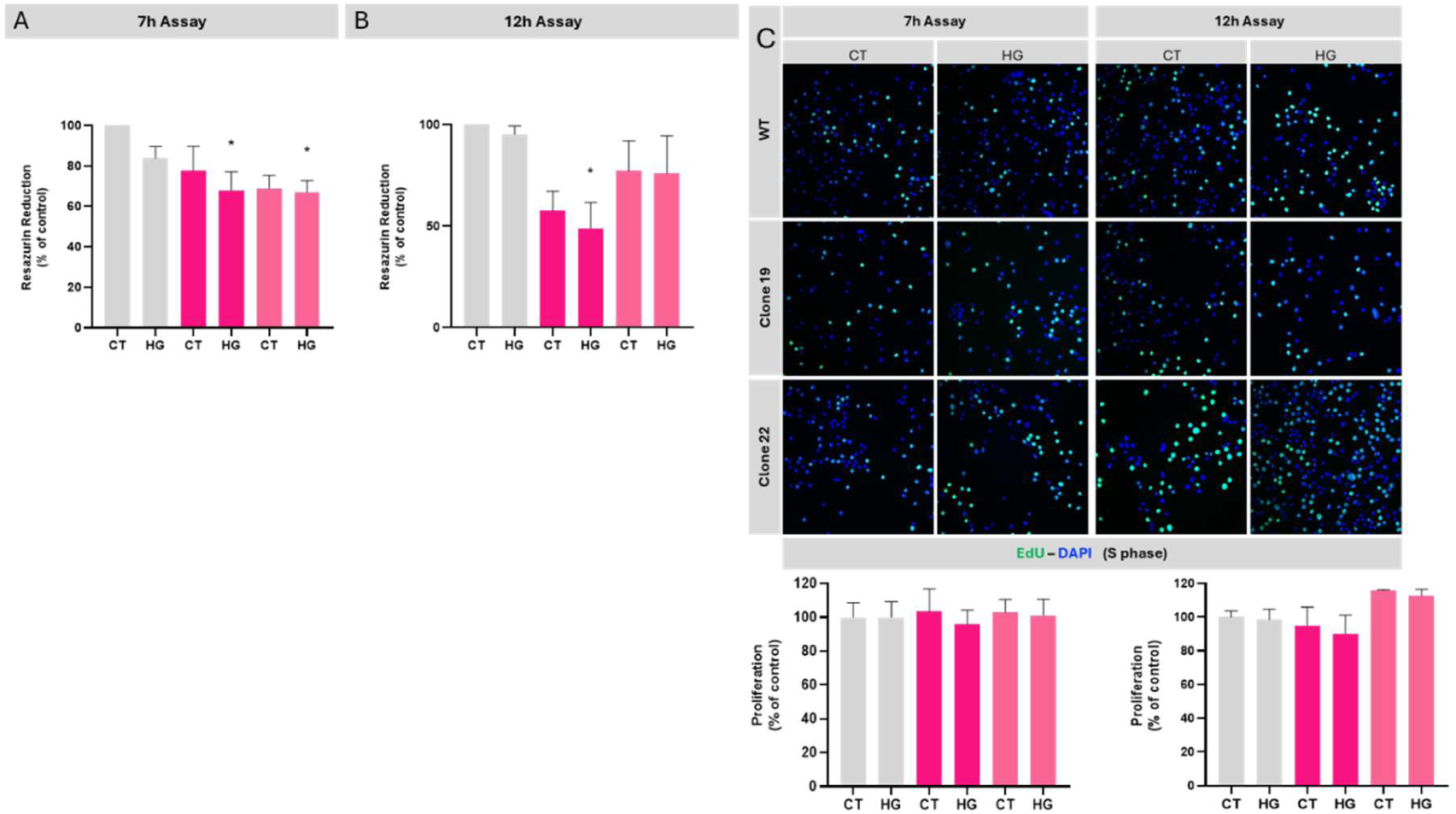
Viability and proliferation of *Ndr2* downregulated BV-2 cultured cells exposed, or not, to high glucose conditions. WT BV-2 (WT) and *Ndr2* downregulated BV-2 (Clone 19 and Clone 22) cell cultures were incubated in 5.5 mM glucose (CT) or 30.5 mM glucose (HG) under different conditions: (A-C) 7h assay; (B-D) 12h assay. The results are presented as the mean ± SEM, and statistical analysis was assessed with a one-way ANOVA test, after confirmation of a Gaussian distribution; * p ≤ 0.05 compared with WT CT; n=6-8 for Alamar Blue assay, and n=5-7 for proliferation assay. EdU (green) incorporation indicates DNA synthesis during cell proliferation and cells were stained with the nuclear marker DAPI (blue).

**Figure 6.**
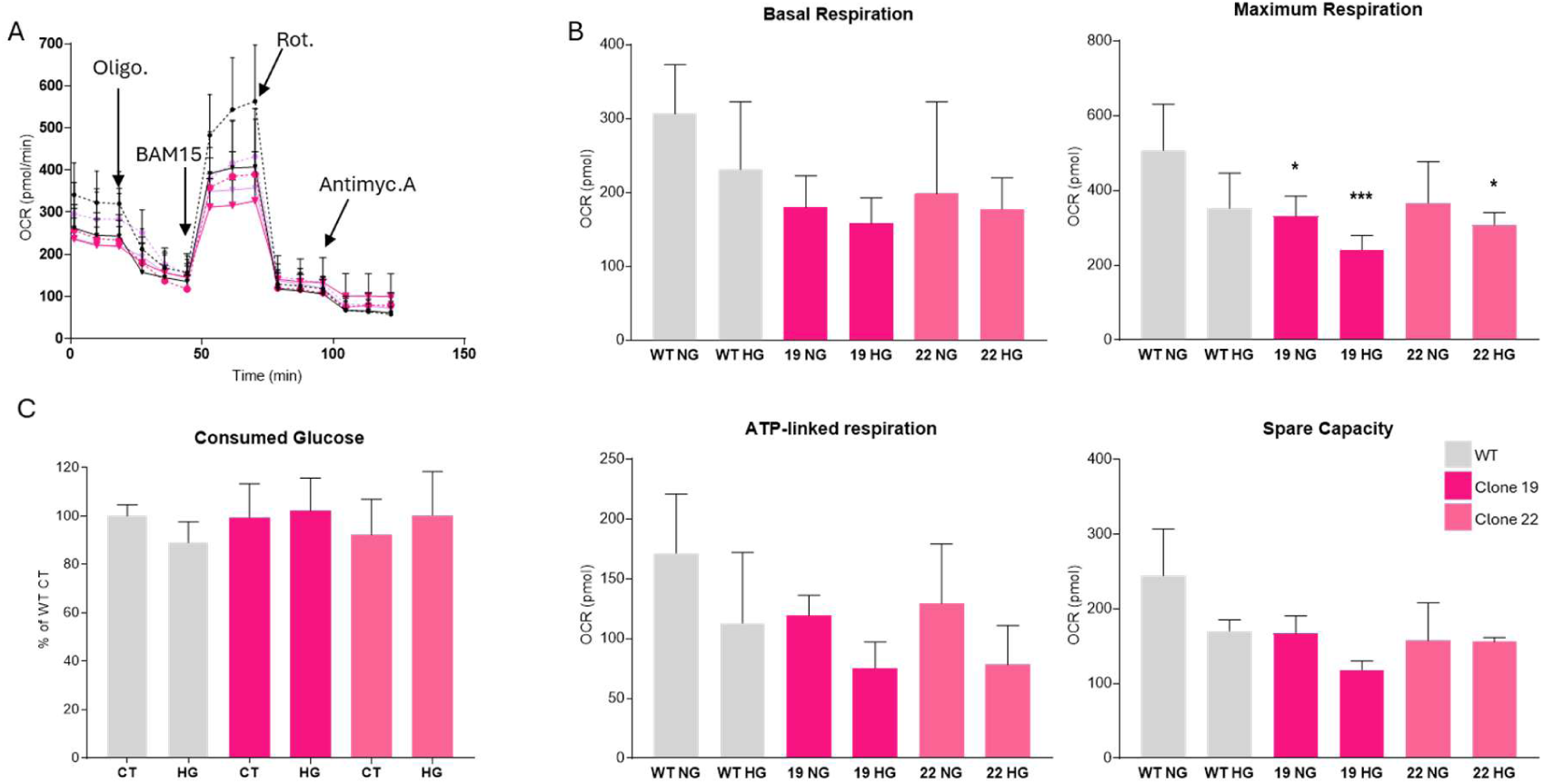
Metabolism of *Ndr2* downregulated BV-2 cells. Effect of *Ndr2* downregulation on (A, B) mitochondrial oxygen consumption measured by the Seahorse assay and on (C) relative glucose consumption measured by glucose test-strip for diabetes from BV-2 microglial cells media after 7h exposure to DMEM containing 5.5mM (NG) or 30.5 mM glucose (HG). The results are presented as the mean ± SEM, and statistical analysis was assessed with a two-way ANOVA test with Turkey correction; * p ≤ 0.05, compared with WT CT; *** p ≤ 0.001, compared with WT CT; n=6-8 independent experiments.

While the resazurin (Alamar Blue) assay provided an indirect assessment of cell viability and metabolic activity, we sought to directly evaluate cell death in WT, clone 19, and clone 22 BV- 2 cells under both control (CT) and high glucose (HG) conditions. To this end, we performed flow cytometry analysis. We evaluated the FITC-annexin-V (AV) and 7-amino-actinomycin D (7-AAD) staining and grouped them into living cells (AV-/7-AAD-), early apoptotic cells (AV+/7- AAD-), late apoptotic cells (AV+/7-AAD+), and necrotic cells (AV-/7-AAD+), as shown in Supp. Figure S3. In CT conditions, the clones 19 and 22 present a similar percentage of live cells than WT BV-2 (WT: 84.5 ± 6.9% of the cells, Clone 19: 83.6± 4.9%, Clone 22: 95.3 ± 0.6%) while they present lower percentages of early apoptotic (WT: 1.1 ± 0.2%, Clone 19: 1.3 ± 0.4%, Clone 22: 1.0 ± 0.1%), late apoptotic/necrotic (WT: 7.5 ± 3.2%, Clone 19: 8.0 ± 2.5%, Clone 22: 2.0 ± 0.3%) and necrotic cells (WT: 5.5 ± 2.4%, Clone 19: 7.1± 2.5%, Clone 22: 2.3 ± 0.7%) than WT BV-2. In the presence of HG conditions, both clones 19 and 22 present a similar, even slightly increased percentage of live cells than WT BV-2 (WT: 78.1 ± 9.7%, Clone 19: 81.1 ± 8.0%, Clone 22: 88.9 ± 4.2%). Similarly to CT conditions, the clones present lower percentages of early apoptotic (WT: 3.6 ± 2%, Clone 19: 1.4 ± 0.4%, Clone 22: 2.5 ± 1.1%), late apoptotic/necrotic (WT: 7.6 ± 2.8%, Clone 19: 9.4 ± 4.1%, Clone 22: 5.4 ± 2.5%) and necrotic cells than WT BV-2 (WT: 10.6 ± 5.3%, Clone 19: 7.1 ± 2.5%, Clone 22: 2.9 ± 1.1%).

While the resazurin (Alamar Blue) assay provides an indirect measure of cell viability and metabolic activity, it does not directly assess cell proliferation. To specifically evaluate proliferative activity, we therefore performed an EdU incorporation assay. No significant alterations were observed either for CT or HG conditions for clones 19 and 22 in both 7h or 12h assays (WT 7h HG: 99.9 ± 9.4% of WT CT; WT 12h HG: 98.5 ± 6.2% of WT 7h CT), (Clone 19 7h CT: 103.2 ± 13.5% of WT 7h CT; Clone 19 7h HG: 96.1 ± 8.2% of WT 7h CT; Clone 19 12h CT: 94.9 ± 10.9% of WT 12h CT; Clone 19 12h HG: 90.0 ± 11.1% of WT 12h CT), (Clone 22 7h CT: 102.8 ± 7.7% of WT 7h CT; Clone 22 7h HG: 101.1 ± 9.5% of WT 7h CT; Clone 22 12h CT: 115.8 ± 0.4% of WT 12h CT; Clone 22 12h HG: 112.3 ± 4.1% of WT 12h CT).

### *Ndr2* downregulation affects the metabolism of microglial cells

The proliferation and apoptosis assays suggest that the decrease in the resazurin reduction by the *Ndr2* downregulated BV-2 compared to the WT BV-2 is due to metabolic changes. This led us to analyze the mitochondrial function using the Seahorse assay (Fig. 5 and Supp. Fig. S4).

We observed a decrease in oxygen consumption rate (OCR) profile at basal level comparing the clones with the WT BV-2 cells, with a lower basal respiration (Clone 19 NG: 74.5 ± 9.8% of CT; Clone 19 HG: 69.2 ± 15.5% of CT; Clone 22 NG: 87.3 ± 14.0% of CT and Clone 22 HG: 68.5 ± 9.3% of CT), as well as maximal respiration (Clone 19 NG: 48.0 ± 20.0% of CT; Clone 19 HG: 52.7 ± 32.1% of CT; Clone 22 NG: 86.9 ± 26.2% of CT and Clone 22 HG: 80.0 ± 3.0% of CT), ATP-linked respiration (Clone 19 NG: 69.0 ± 9.0% of CT; Clone 19 HG: 43.9 ± 12.8% of CT; Clone 22 NG: 75.4 ± 29.0% of CT and Clone 22 HG: 45.0 ± 19.0% of CT) and spare respiratory capacity (Clone 19 NG: 68.0 ± 9.0% of CT; Clone 19 HG: 48.3 ± 5.0% of CT; Clone 22 NG: 64.5 ± 20.7% of CT and Clone 22 HG: 63.0 ± 2.0% of CT). We observed the expected decrease in OCR compared to WT HG with WT NG cells (basal respiration WT HG: 76.4 ± 20.8% of CT; maximal respiration WT HG: 75.8 ± 27.7% of CT; ATP-linked respiration WT HG: 65.9 ± 34.4% of CT; spare respiratory capacity WT HG: 69.7 ± 6.2% of CT) (Fig 5.A-B). Moreover, we measured the glucose consumed by the WT and *Ndr2* clones BV-2 in control and HG conditions. These results demonstrate that HG negatively affects microglial cells even during acute exposures. Moreover, contrary to WT BV-2 cells, in which we observed a significant difference of glucose consumption between HG and CT conditions, the Nr2 downregulated BV-2 cells did not consume more glucose when exposed to a higher glycemic condition (Fig. 5C). These results suggest that *Ndr2* downregulation affects the metabolism of microglial cells but does not impair their proliferation, even in HG conditions.

### High glucose and *Ndr2* downregulation affect phagocytic efficiency

To further define how *Ndr2* downregulation affects important functions of microglial cells, we investigated the phagocytic capacity of WT and *Ndr2*downregulated BV-2 cells, in HG and CT conditions, using fluorescent yellow-green latex beads (Figure 5). WT BV-2 cells present a basal phagocytosis efficiency of 115.1 ± 18.7% (7h assay) and 149.7 ± 8.3% (12h assay) while the WT BV-2 cells present a phagocytosis efficiency of 105.6 ± 9.7 % when exposed to HG for 7h and of 106.2 ± 10.1% when exposed to variation of glucose. The *Ndr2* downregulated BV- 2 cells exposed to NG conditions for 7h show a non-statistically significant decrease of phagocytic efficiency compared to WT BV-2 cells (Clone 19 7h CT: 70.6 ± 21.6%; Clone 22 7h CT: 81.1 ± 17.8%) and HG conditions (Clone 19 7h HG: 47.2 ± 2.3%; Clone 22 7h HG: 87.1 ± 18.7%). For the 12h assay, a significant decrease was observed between *Ndr2* downregulated BV-2 cells and WT cells, for both clones (Clone 19 12h CT: 46.3 ± 18.9%; Clone 22 12h CT: 100.8 ± 12.1%; Clone 19 12h HG: 38.0 ± 8.7%; Clone 22 12h HG: 95.8 ± 15.8%). Interestingly, the basal phagocytic efficiency of the WT BV-2 CT cells in the 12h assay is higher than in the 7h assay (WT 7h CT: 115.05 ± 18.7%; WT 12h CT: 149.7 ± 8.3%) suggesting the reactivity of the BV-2 cell cultures to the change of medium without alteration of the glucose levels. The basal phagocytic efficiency of WT HG is similar in both the 7h and the 12h assays (WT 7h HG: 105.6 ± 9.7%; WT 12h HG: 106.2 ± 10.1%).

### High glucose and *Ndr2* downregulation decrease BV-2 migration

To assess the impact of *Ndr2* downregulation on BV-2 microglia migration in CT and HG conditions, we performed the Boyden Chamber assay using transwell cell culture inserts with 8.0 µm pore diameter. WT BV-2 cells and *Ndr2* downregulated BV-2 clonal populations were exposed to NG or HG for 7h, and migrating cells, located on the bottom side of the insert, were fixed and stained with DAPI to allow cell counting. *Ndr2* downregulated BV-2 cells show a decreased migration compared to the WT parental cells, in HG conditions (Clone 19 7h CT: 83.7 ± 17.6% of WT CT; Clone 19 7h HG: 55.9 ± 18.3% of WT CT; Clone 22 7h CT: 55.8 ± 19.2% of WT CT; Clone 22 7h HG: 25.6 ± 5.2% of WT CT). Interestingly, the migration ability of the BV-2 cells exhibited a non-statistically significant decrease in cell migration in HG conditions (WT 7h HG: 65.4 ± 9.0 % of WT CT). We observed a similar effect of HG on BV-2 cells migration in *Ndr2* downregulated BV-2 cells (Figure 7).

**Figure 7.**
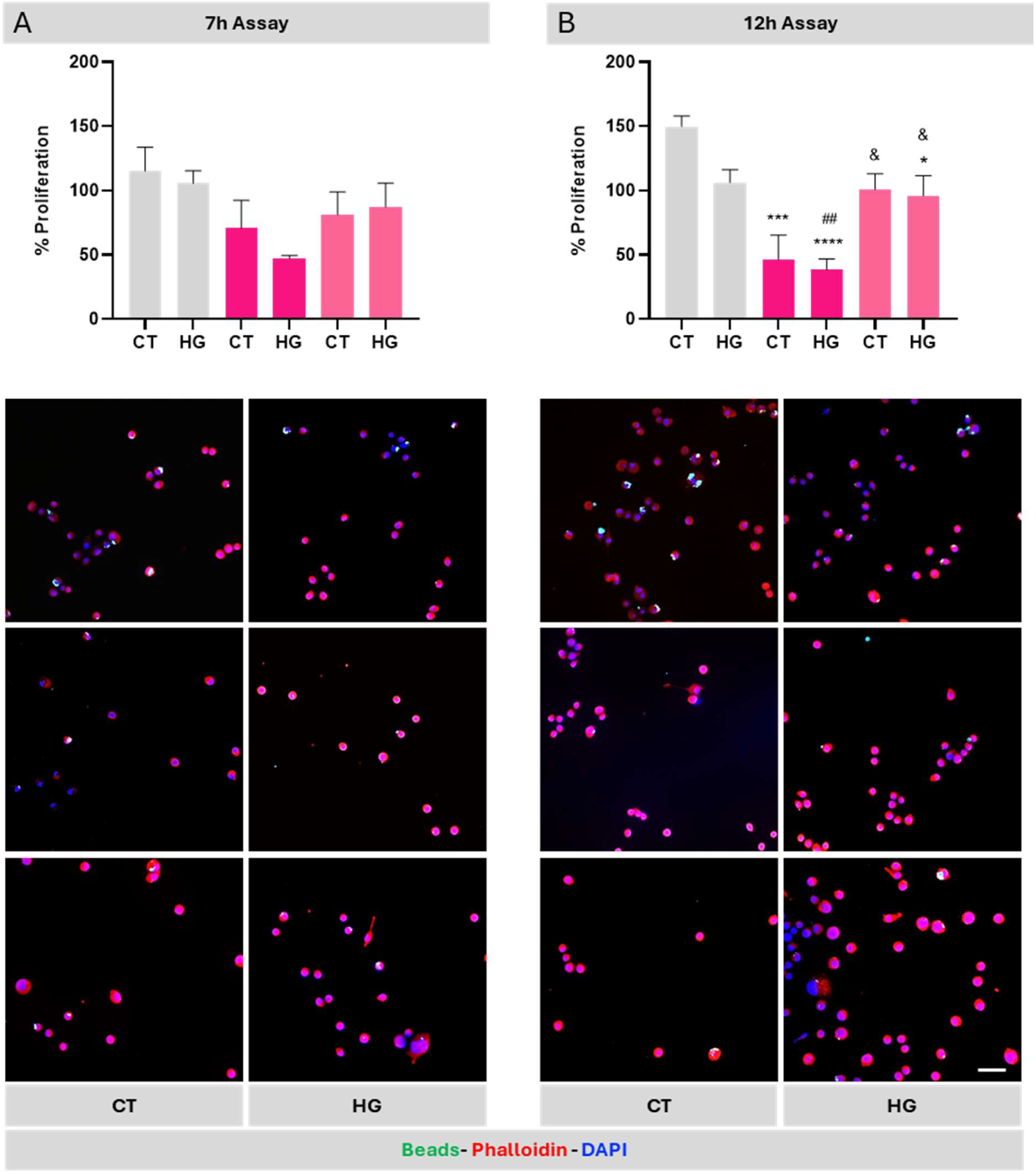
***Ndr2* downregulation decreases BV-2 phagocytic efficiency**. WT BV-2 (WT) and *Ndr2* downregulated BV-2 (Clone 19 and Clone 22) cell cultures were incubated in 5.5 mM glucose (CT) or 30.5 mM glucose (HG) for different periods: (A) 7h assay; (B) 12h assay. The results are presented as the mean ± SEM, and statistical analysis was assessed with a one-way ANOVA test after confirmation of a Gaussian distribution. * p ≤ 0.05, *** p ≤ 0.001, **** p ≤ 0.0001, compared with WT CT; ## p ≤ 0.001, compared with WT HG; & p ≤ 0.05, compared with Clone 19 HG; n=4-6. Fluorescent beads (green) were used to assess the phagocytic activity, and cells were stained with phalloidin (red) and the nuclear marker DAPI (blue). Scale bar: 50 µm.

### *Ndr2* downregulation and exposure to HG or LPS upregulate the expression of IL-17a and TNF

To determine the role of NDR2 kinase in cytokine’s secretion by microglial cells, we compared the profile of secreted cytokines by WT and *Ndr2* downregulated cells (clone 19) exposed or not to HG for 7h, using semi-quantitative antibody microarrays. The levels of various cytokines and chemokines, such as IL-10, IL-6, thrombopoietin (TPO), IL-17a, TNF, increased in the cell medium of clone 19 exposed to NG (19-CM CT) and exposed to HG (19-CM HG) compared with WT-CM CT and WT-CM HG, respectively. Two pro-inflammatory factors, the chemokine MCP5 and the receptor 1 for the pro-inflammatory cytokine TNF-alpha (TNFRI) were significantly decreased in the cell medium of clone 19 CT and HG compared to the cell medium of WT CT and HG, respectively (Figure 8A).

**Figure 8.**
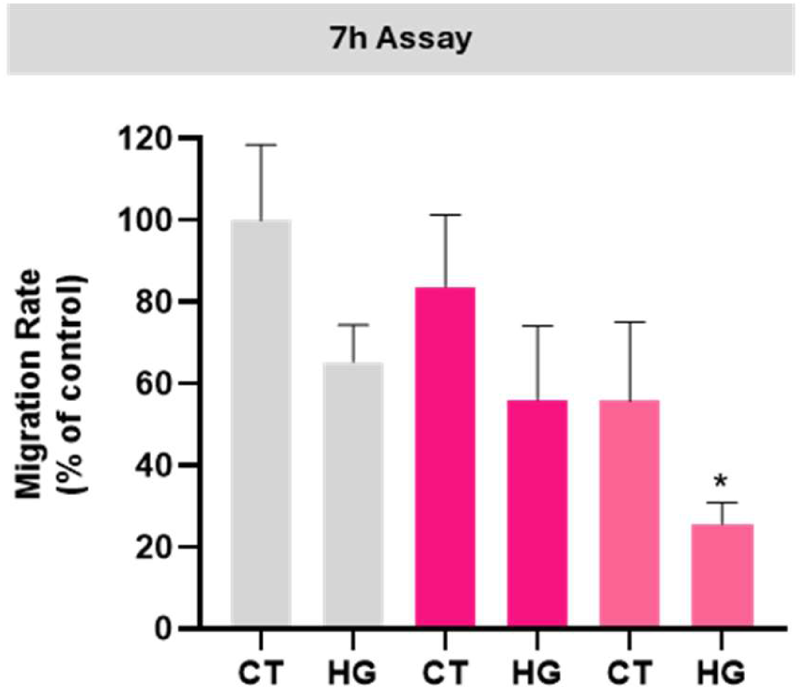
**Migration rate of WT and *Ndr2* downregulated BV-2 cultured cells exposed or not to high glucose**. WT BV-2 (WT) and *Ndr2* downregulated BV-2 (Clone 19 and Clone 22) cell cultures were incubated in 5.5 mM glucose (CT) or 30.5 mM glucose (HG) for 7h. The results are presented as the mean ± SEM, and statistical analysis was assessed with a one-way ANOVA test after confirmation of a Gaussian distribution; * p ≤ 0.05, compared with WT CT (n= 4-7).

**Figure 9.**
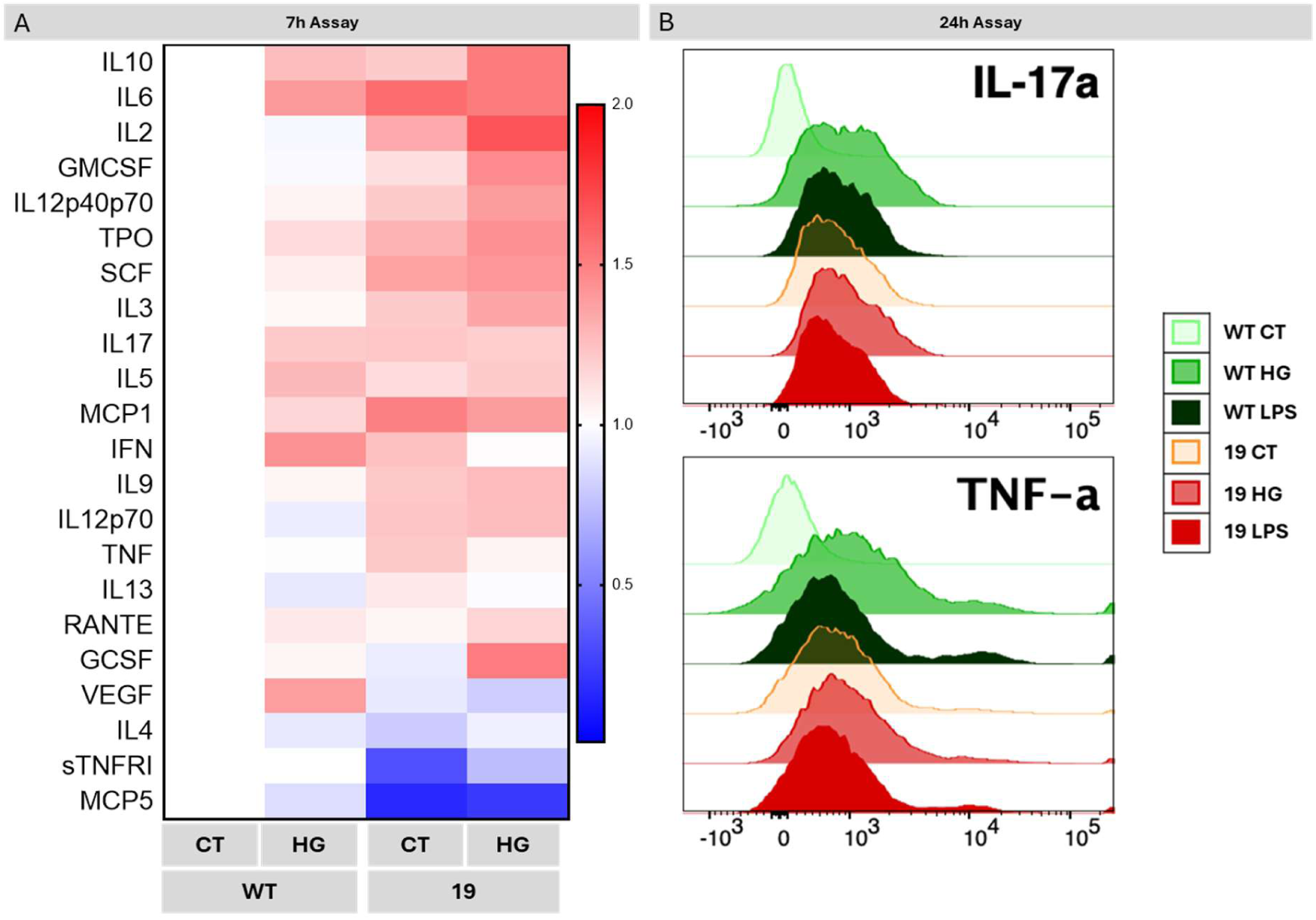
Expression of cytokines/chemokines in WT and *Ndr2* downregulated BV-2 cells exposed to HG. (A) Cytokine array analysis demonstrates prominent upregulation of various cytokines in the conditioned medium (CM) of *Ndr2* downregulated BV-2 cells. BV-2 cell cultures were incubated with 5.5 mM glucose (CT) or 30.5 mM glucose (HG) for 7h. (B) IL-17a and TNF levels were evaluated by flow cytometry analysis. BV-2 cell cultures were incubated in 5.5 mM glucose (CT), 30.5 mM glucose (HG) or 100 ng/mL LPS (LPS) for 24h. The images of the different membranes of the cytokine arrays and their map are represented in Supp. Fig. S6.

These results were consolidated by the assessment of the expression of TNF and IL-17 after 24h exposure to HG or LPS (positive control, 100 ng/mL) by flow cytometry. For the analysis, CD11b-V500 and P2γ12-APC positive cells were selected, which correspond to microglial cells, and then the expression of IL-17a and TNF was evaluated. As we can see in Figure 8B, in the case of WT BV-2 cells, HG promoted the expression of IL-17a and TNF, when compared to the control condition. Similarly, the *Ndr2* downregulation led to an increased expression of IL-17a and TNF to similar levels as LPS, and exposure to HG did not further alter their expression levels in comparison to the control. This pattern is consistently observed in response to LPS exposure as well. In conclusion, *Ndr2* downregulation is sufficient to induce an increase in pro-inflammatory cytokines production and secretion by microglial cells, regardless of the glycose concentration in the cell culture media.

## Discussion

Hyperglycemic environments are known to disrupt cellular metabolism and activate signaling pathways such as the Hippo pathway, which modulates the expression and stability of NDR kinases[5, 9, 32]. NDR1 and NDR2, key effectors of the Hippo pathway, are implicated in regulating cell proliferation, apoptosis, differentiation, and immune responses, including innate and antiviral defenses [6, 8, 10, 12, 17, 33]. Our study demonstrates that NDR2 is a crucial regulator of microglial metabolic adaptation and immune functionality under hyperglycemic stress. We found that NDR2 protein levels increase in BV-2 microglial cells following high glucose (HG) exposure and glycemic fluctuation. We also detected elevated NDR1/2 protein expression following 12-hour glycemic fluctuation (data not shown). These findings suggest a role for NDR kinases in the microglial response to metabolic stress. Moreover, we demonstrated that *Ndr2* downregulation impairs mitochondrial respiration, metabolic flexibility, phagocytosis, migration, and cytokine homeostasis, indicating a multifaceted role for NDR2 in maintaining microglial resilience and function in a metabolically challenging environment.

Previous studies have implicated NDR kinases in modulating inflammation[8, 10, 17]. We hypothesized that the increased NDR2 expression observed under HG conditions may be part of a compensatory inflammatory response to restore homeostasis. However, no significant change in *Ndr2* mRNA levels was observed under the same conditions, suggesting that regulation occurs at the post-transcriptional level. This may involve enhanced protein translation, increased stability, or reduced degradation under hyperglycemic stress. Further studies are warranted to clarify the mechanisms of this regulation.

To explore the functional role of NDR2 in microglial response to glucose stress, we generated *Ndr2* downregulated BV-2 clones via CRISPR-Cas9 targeting exon 7 of the *Ndr2/Stk38l* gene. Although we did not achieve complete knockout, these cells exhibited distinct phenotypes. Viability assays using resazurin showed reduced metabolic activity in *Ndr2* downregulated clones after 7 hours of HG exposure, compared to wild-type (WT) cells under control conditions. This reduction was not due to increased cell death, as confirmed by apoptosis and necrosis assays, nor due to altered cell cycle progression, as shown by EdU incorporation. These findings suggest that NDR2 is critical for sustaining cellular metabolism under stress, rather than directly regulating survival or proliferation.

Short-term HG exposure did not affect WT microglial viability, indicating a degree of resilience. However, prolonged HG exposure increased oxidative and nitrosative stress, leading to elevated rates of apoptosis and necrosis. These findings are consistent with literature showing that the extent and duration of hyperglycemia dictate microglial stress responses[31, 34].

Seahorse analysis further revealed a significant decrease in basal oxygen consumption rate (OCR) in *Ndr2* downregulated clones, along with impaired mitochondrial respiration, including reduced maximal respiration, proton leak, and spare respiratory capacity. Despite unaffected viability, these cells displayed lower metabolic activity, reinforcing the role of NDR2 in maintaining mitochondrial function and oxidative phosphorylation under normal and HG conditions.

Strikingly, while WT cells adapted to HG by reducing OCR and shifting toward glycolysis—a known metabolic reprogramming response — *Ndr2* downregulated cells failed to do so. This indicates a defect in metabolic flexibility and suggests a role for NDR2 in orchestrating energy pathway transitions. Moreover, increased proton leak and elevated complex III activity in *Ndr2* downregulated cells under HG conditions point to mitochondrial dysfunction and potential compensatory mechanisms to sustain ATP production. The absence of adaptive glucose consumption changes in these cells further supports impaired metabolic sensing and regulation.

Beyond energy metabolism, NDR2 appears to influence key effector functions of microglia. NDR2 colocalized with IBA1, an actin-crosslinking protein critical for membrane ruffling and phagocytosis. Localization of NDR2 to membrane ruffles and process tips suggests a role in cytoskeletal remodeling. Consistently, *Ndr2* downregulated cells exhibited reduced phagocytic activity under both control and HG conditions, likely due to a combination of energy deficits and actin cytoskeletal dysfunction.

Microglial migration, assessed using a Boyden chamber assay, was similarly impaired in *Ndr2* downregulated cells, again under both basal and HG conditions. Migration relies on ATP-dependent actin remodeling and purinergic signaling, and our findings support a role for NDR2 in regulating this function, likely through effects on focal adhesion and actin dynamics. Functionally, these impairments in phagocytosis and migration were time- and glucose-dependent. Phagocytosis was significantly reduced under 7-hour and 12-hour HG exposure in *Ndr2* downregulated cells, whereas WT cells maintained or even enhanced phagocytic capacity under similar conditions. This underscores the role of NDR2 in facilitating adaptive responses to metabolic stress and maintaining microglial functionality.

Importantly, although *Ndr2* downregulated cells remained viable and proliferative, their impaired functional outputs suggest that NDR2 selectively regulates energy-dependent immune functions rather than general cell survival.

Microglia are metabolically flexible cells, typically relying on oxidative phosphorylation under homeostasis[20, 35]. However, during neuroinflammation or injury, they switch to glycolysis to meet increased energy demands and support inflammatory cytokine production. Inhibiting glycolysis dampens microglial activation, further linking metabolism to immune function[36–38].

Consistent with this, we found that *Ndr2* downregulation induced a dysregulated inflammatory phenotype, particularly under HG conditions. *Ndr2* downregulated clones exhibited increased secretion of pro-inflammatory cytokines including TNF, IL-6, and IL-17, even without LPS stimulation. Interestingly, some cytokines such as RANTES, MCP-5, and sTNFRI were downregulated, suggesting an imbalanced or mixed activation profile. The decrease in sTNFRI may reduce the buffering of TNF signaling, potentially enhancing cytotoxic effects. Meanwhile, upregulation of IL-10 may reflect an attempted compensatory resolution of inflammation[39–42].

This complex, dysregulated cytokine profile is characteristic of microglial responses under chronic inflammatory conditions and may reflect aberrant activation of pathways such as NF- κB or MAPK.[40] Thus, NDR2 appears to be a central modulator of microglial inflammatory homeostasis. Its downregulation mirrors feature seen in chronic neuroinflammatory states, including those observed in diabetic retinopathy, Parkinson’s disease, Alzheimer’s disease, and autoimmune neurodegeneration [48].

A major strength of this study is the use of physiologically relevant glucose exposure time points—specifically 7-hour and 12-hour conditions—to model both acute hyperglycemia and stress induced by daily glycemic variability. This approach more accurately reflects the transient but impactful glucose fluctuations observed in individuals with diabetes and metabolic disorders, thereby enhancing the translational value of our findings.

Nonetheless, our study has several limitations. The use of partial *Ndr2* downregulation, rather than a complete knockout, may limit the interpretation of the full functional spectrum of NDR2. Additionally, although BV-2 cells are a widely accepted model for microglia, they may not fully recapitulate the behavior of primary microglia within the central nervous system microenvironment. It also remains unclear which direct downstream targets of NDR2 are responsible for its regulatory effects on mitochondrial function and cytoskeletal dynamics.

To strengthen translational relevance, *in vivo* studies are needed to validate these findings in a more physiologically and pathologically accurate context. These studies should also examine whether modulating NDR2 can reverse established inflammatory phenotypes or neurodegenerative changes. Furthermore, future investigations should explore whether other components of the Hippo signaling pathway similarly influence microglial metabolism and function, potentially unveiling broader therapeutic targets for neuroinflammatory and metabolic diseases.

## Conclusion

Our findings position NDR2 as a key regulator of microglial mitochondrial function, metabolic flexibility, and effector responses such as phagocytosis and migration. While short-term glycolytic activation supports microglial responses, chronic reliance on glycolysis—especially in the context of impaired mitochondrial function—leads to pro-inflammatory bias, reduced functional plasticity, and diminished debris clearance. *Ndr2* downregulation recapitulates this phenotype, highlighting its role in metabolic adaptation and immune regulation.

Given the prevalence of metabolic dysregulation in neurodegenerative and metabolic diseases, NDR2 represents a promising therapeutic target for restoring microglia homeostasis and mitigating chronic inflammation. Future research should explore the molecular pathways downstream of NDR2 that govern metabolic reprogramming, cytoskeletal dynamics, and cytokine signaling in microglia.

## Permission/acknowledgement

The authors would like to thank the support of iLAB – Microscopy and Bioimaging Lab, a facility of the Faculty of Medicine, University of Coimbra (FMUC), and a member of the national infrastructure PPBI-Portuguese Platform of BioImaging (POCI-01-0145-FEDER- 022122) and supported by FSE CENTRO-04-3559-FSE-000142. We are indebted to numerous colleagues from the University of Coimbra community who have provided insightful advice, language help, and proofreading of this article. In particular, the authors would like to thank Dr. Ana Luisa Cardoso (Centre for Neuroscience and Cell Biology - CNC, UC) for kindly providing the iPSC-derived microglial cells used in this study, Dr. Anton Lennikov (Schepens Eye Research Institute, Harvard Medical School) and Dr. Raquel Santiago (Coimbra Institute for Clinical and Biomedical Research, UC) for their help with setting up primary microglial cell culture as well as Dr. José Teixeira (CNC, UC) for his help with our seahorse assay. We deeply appreciate their expertise and generosity in supporting our research. Finally, the authors would like to thank all the undergraduates who helped during the length of this study.

## Social media

Scientists have uncovered a key player in microglial cell function: Ndr2 kinase. This protein helps microglia adapt to high glucose levels and regulates inflammation in the retina.

## Key findings

● NDR2 is crucial for microglial metabolism under high glucose conditions.
● Downregulating Ndr2 impairs phagocytosis and migration of microglia.
● Lower NDR2 levels lead to increased pro-inflammatory cytokines production.
This breakthrough could lead to new treatments for diabetic retinopathy and other neuroinflammatory conditions. Let’s keep an eye on NDR2 kinase research! #DiabeticRetinopathy #MicroglialCells #MedicalResearch #universidadedecoimbra #befmuc #icbr #fa_lab.uc #saude

## Abbreviations

7-AAD: 7-aminoactinomycin D
ADP: adenosine diphosphate
ATP: adenosine triphosphate
AV: annexin V
CT: control
ECAR: extracellular acidification rate
EdU: 5-ethynyl-2’-deoxyuridine
GFAP: glial fibrillary acidic protein
HG: high glucose
IBA1: ionized calcium binding adapter protein 1
IL-6: interleukin-6
IL-10: interleukin-10
IL-17a: interleukin-17 a
iPSC: induced pluripotent stem cells
KD: knockdown
NDR: nuclear dbf2-related
NG: normal glucose
NO: nitric oxide
OCR: oxygen consumption rate
qRT-PCR: quantitative reverse transcription
PCR ROS: reactive oxygen species
SDS: sodium dodecyl sulfate
SEM: standard error of the mean
TNF: tumor necrosis factor
WT: wild type

---

**Table 1.**
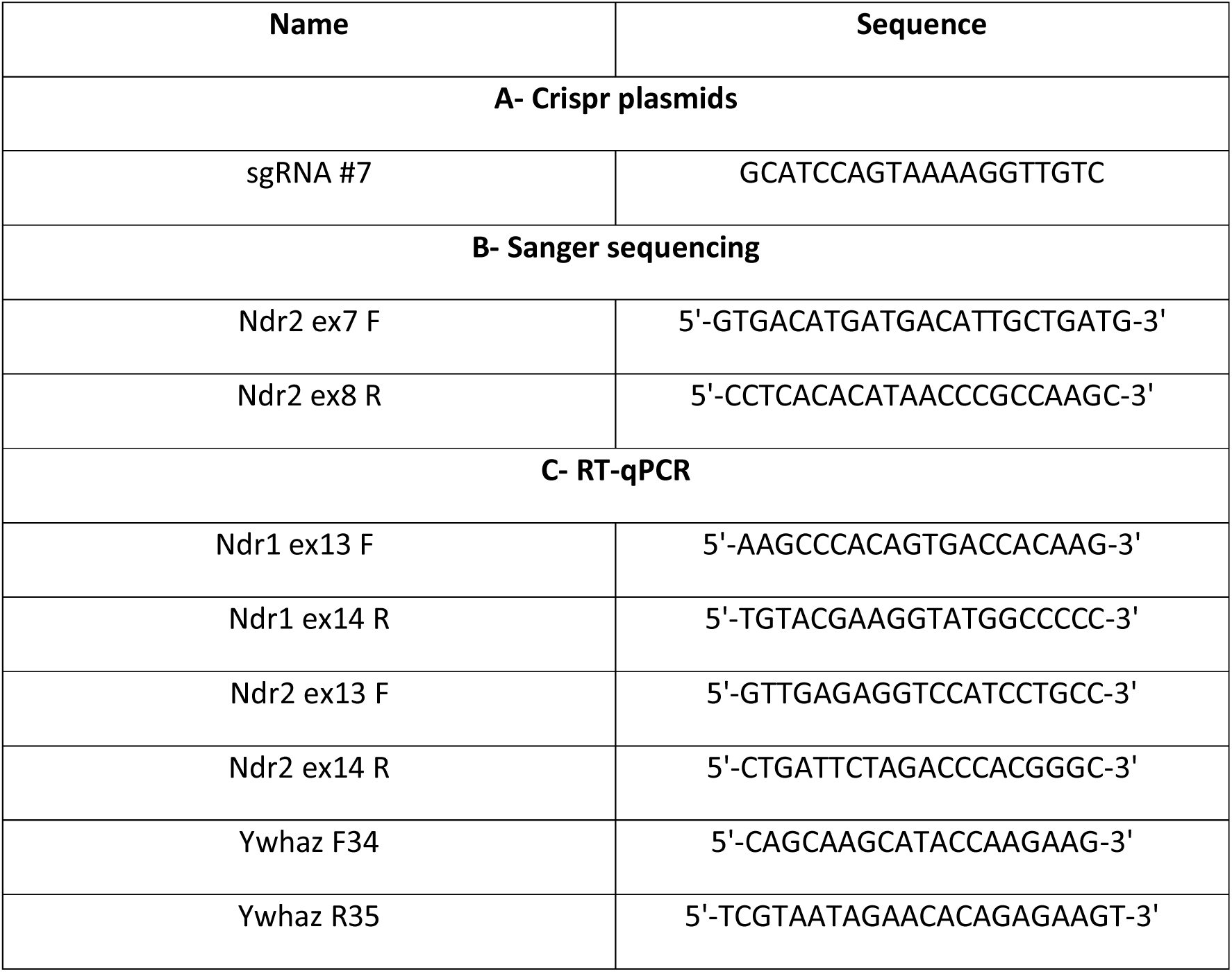
Genotyping primers. . (A) single-guide RNA (sgRNA) sequence for against *Ndr2* exon 7; (B) Oligonucleotide primers for Sanger sequencing BV-2, targeting the *Ndr2* exons 7 and 8; (B) Oligonucleotide primers for RT-qPCR against the mouse *Ywhaz* gene and against the exons 13 and 14 of the mouse *Ndr1* and *Ndr2* genes. All the primers are reported with the symbol of the corresponding gene.

**Table 2.** Antibodies. List of antibodies used for immunocytochemistry (ICC), western blot (WB) and flow cytometry. Antibodies are reported with the symbol of the corresponding protein (antigen), source including commercial company name (catalogue number) and concentrations used for either IHC or WB.

| Primary antibodies |  |  |  |  |  |  |
| --- | --- | --- | --- | --- | --- | --- |
| Antigen | Host | Company | Reference | ICC conc. | WB conc. | Flow cytometry |
| Ndr1/2 (E-2) | Mouse monoclonal | Santa Cruz | sc-271703. | 1/100 | 1/1000 | - |
| Ndr2 | Rabbit polyclonal | St John's laboratory | STJ94368 | 1/100 | 1/1000 | - |
| Iba1 | Rabbit polyclonal | Wako | 019-19741 | 1/500 | - | - |
| GFAP | Chicken polyclonal | Millipore | AV5541 | 1/100 | - | - |
| Calnexin | Goat polyclonal | Sicgen | AB0041-500 | - | 1/1000 | - |
| V450-A::IL-17a |  |  |  | - | - | 1x |
| PE-A::TNFa |  |  |  | - | - | 1x |
| Annexin V |  |  |  | - | - | 1x |
| 7-AAD |  |  |  | - | - | 1x |
| Phalloidin Tritc Labeled Mixed Isomers |  | Sigma | P1951 | 1x | - | - |
| **Secondary antibodies** |  |  |  |  |  |  |
| **Antigen** | **Host** | **Company** | **Reference** | **ICC conc.** | **WB conc.** | **Fluorophore** |
| anti-Mouse IgG (H+L) | Goat polyclonal | ThermoFisher Sc. | A-11004 | 1/500 | - | Alexa 568 |
| anti-Rabbit IgG (H+L) | Goat polyclonal | Invitrogen | A11008 | 1/500 | - | Alexa 488 |
| anti-Chicken IgG (H+L) | Goat polyclonal | Invitrogen | A11041 | 1/500 | - | Alexa 568 |
| StarBright Blue 700 Goat Anti-Rabbit IgG | Goat polyclonal | Bio-Rad | 12004162 | - | 1/10 000 | Blue 700 |
| HRP Anti-Goat | Rabbit polyclonal | Thermo Fisher Scientific | LTI 611620 | - | 1/10 000 | - |

**Table 3.**
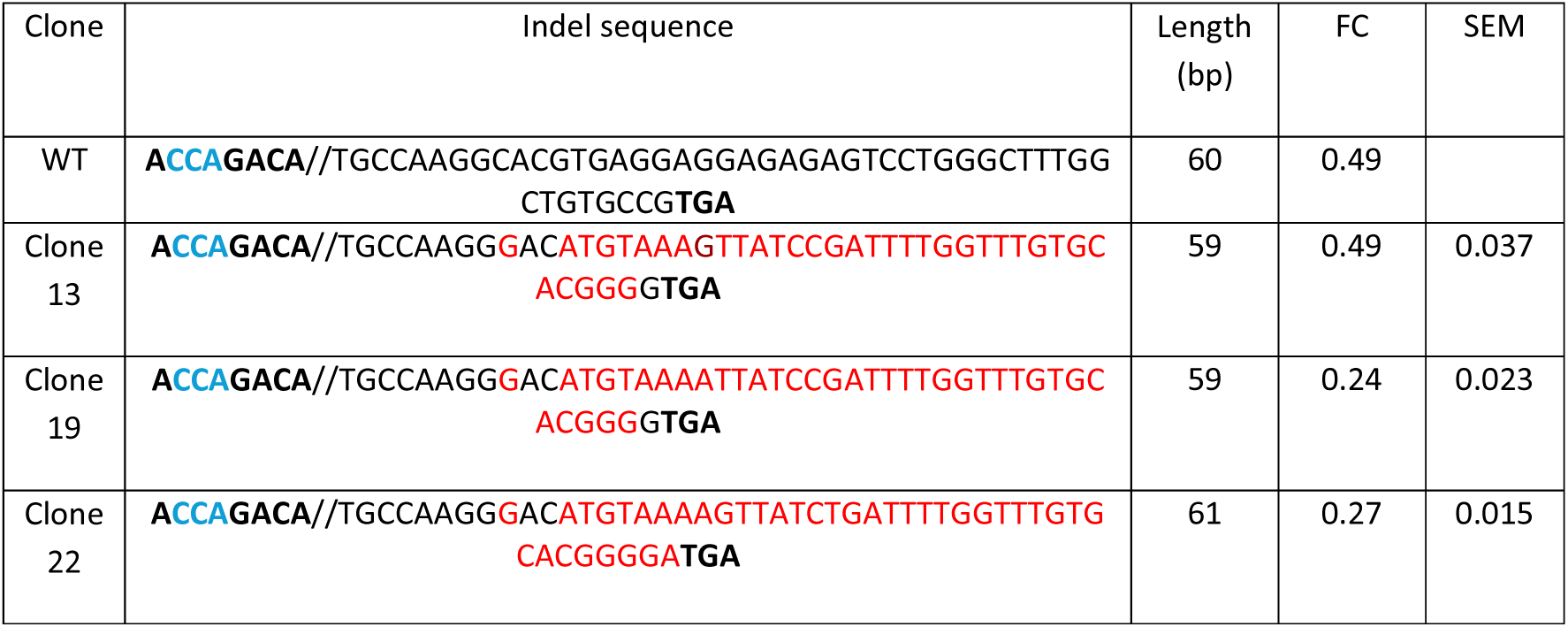
Ndr2 Indel sequences. Three independent *Ndr2/Stk38l* alleles (Clone 13, Clone 19 and Clone 22) generated from independent clonal BV-2 populations, were analyzed by DNA sequencing. The DNA sequence of WT and the clones are presented. Clone 13 contains a mutation in *Ndr2* exon 7 (red) that yields a minus 1 bp frame shift mutation and a downregulation of Ndr2 expression (FC= 0.49 ± 0.07). Clone 19 contains a mutation in *Ndr2* exon 7 (red) that yields a minus 1 bp frame shift mutation and a downregulation of Ndr2 expression (FC= 0.24± 0.05). Clone 22 contains a mutation in *Ndr2* exon 7 (red) that yields a minus 1 bp frame shift mutation and a downregulation of Ndr2 expression (FC= 0.27± 0.03). The results are expressed as fold change (FC) ± SEM (n=2). The PAM sequence is represented in blue, the WT sequence in black and the mutated sequence in red.

## Supplementary Figures

**Supp. Figure S1.**
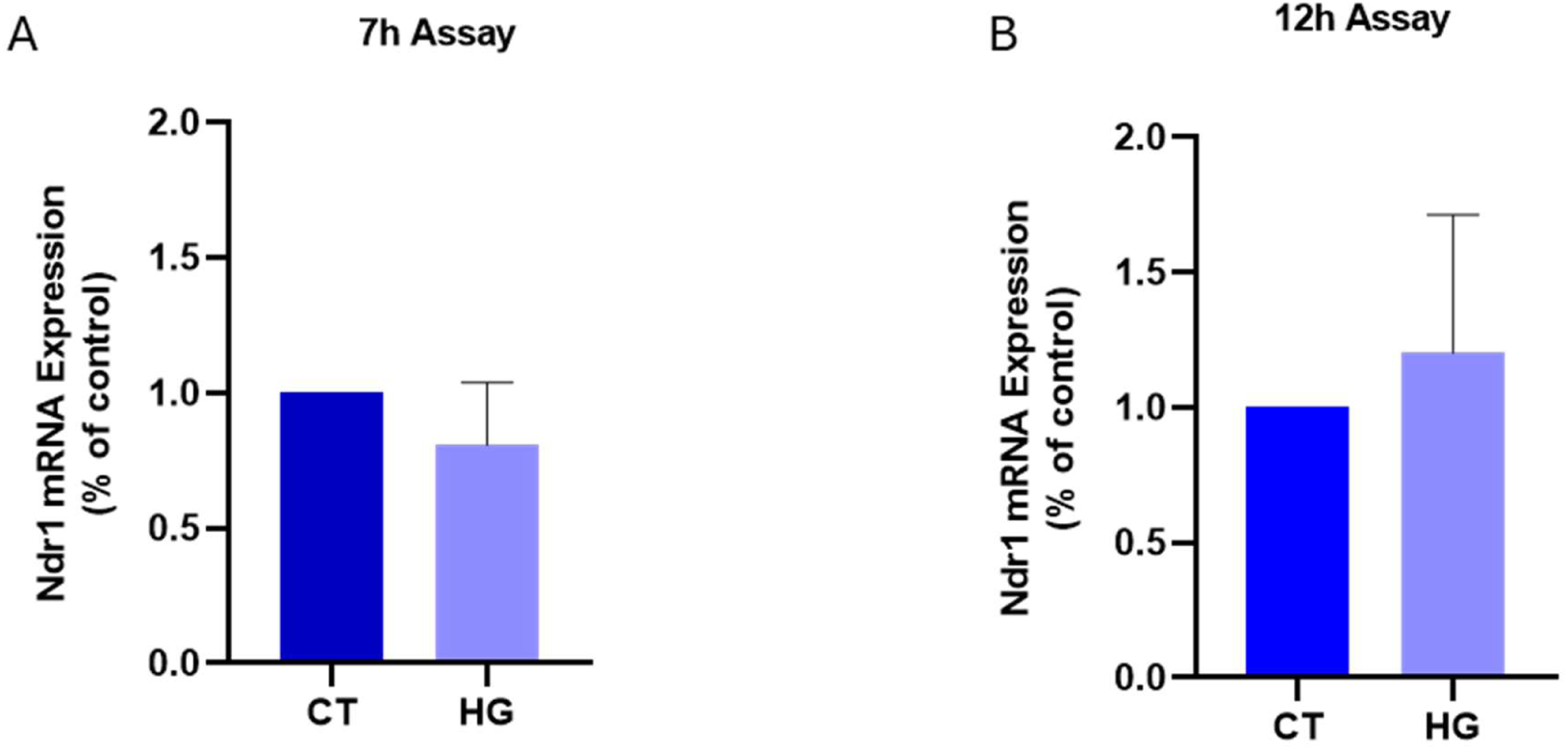
High glucose does not affect the *Ndr1* mRNA expression in BV-2 microglial cells. Expressed. mRNA levels of *Ndr1* in BV-2 cells exposed to 5.5 mM glucose (CT) or 30.5 mM glucose (HG) in both the 7h assay and the 12h assay were evaluated by qRT-PCR analysis. The results are normalized to control and are expressed as mean ± SEM. Statistical analysis was assessed with Student’s t-test; n=3-4 independent experiments.

**Supp. Figure S2.**
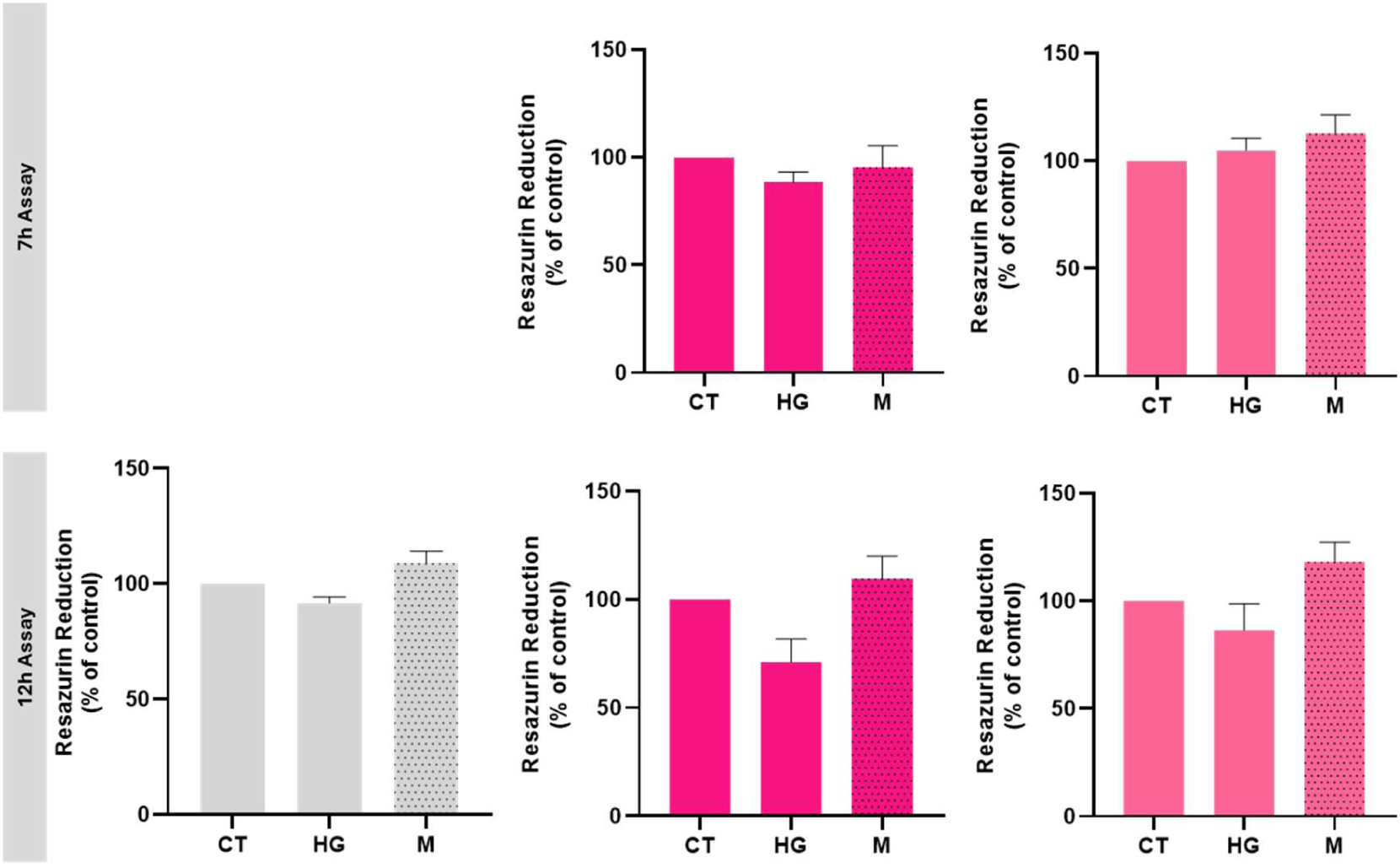
Viability assessment of cultured BV-2 cells exposed to high glucose or mannitol (osmotic control). WT BV-2 (WT) and *Ndr2* downregulated BV-2 (Clone 19 and Clone 22) cell cultures were incubated in 5.5 mM glucose (CT), 30.5 mM glucose (HG) or 25 mM mannitol (M), for different periods: (A) 7h assay; (B) 12h assay. Viability was assessed using the Alamar assay. The results are presented as mean ± SEM. Statistical analysis was assessed with a one-way ANOVA test after confirmation of a Gaussian distribution; * p ≤ 0.05, compared with WT CT; n=6-8 independent experiments.

**Supp. Figure S3.**
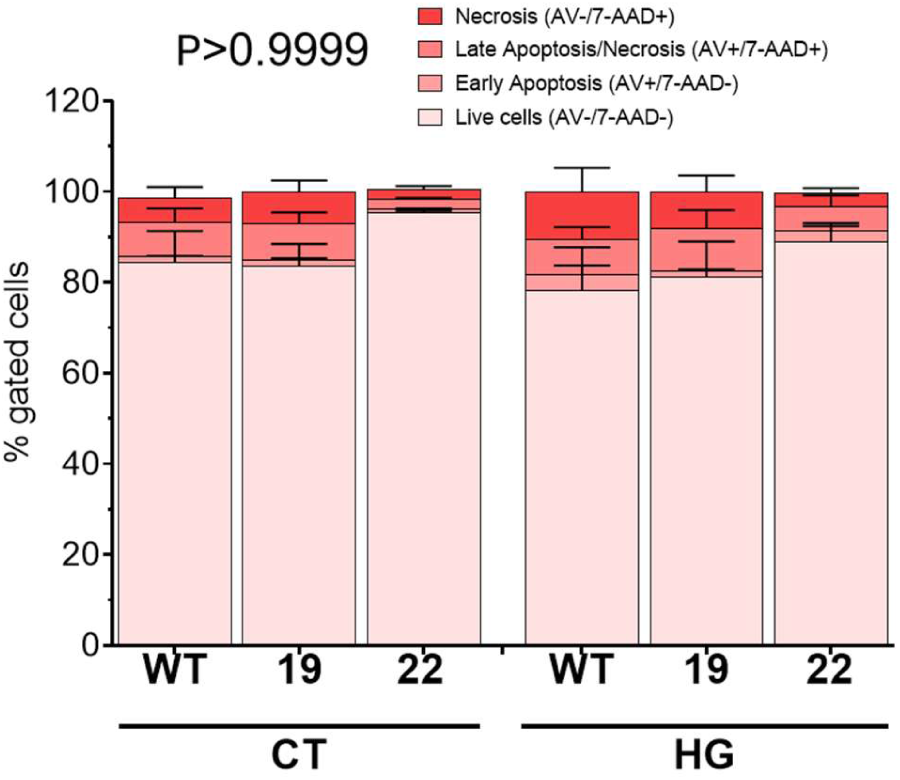
Determination of apoptotic cell fractions of WT and *Ndr2* downregulated BV-2 cells under control and high glucose conditions. Living cells (ANN-V-/7-AAD-), early apoptotic cells (ANN-V+/7-AAD-), late apoptotic/necrotic cells (ANN-V+/7-AAD+) and necrotic cells (ANN-V-/7-AAD+). Asterisks indicate significances compared to control (* p < 0.05; ** p < 0.01); n=3-4 independent experiments.

**Supp. Figure S4.**
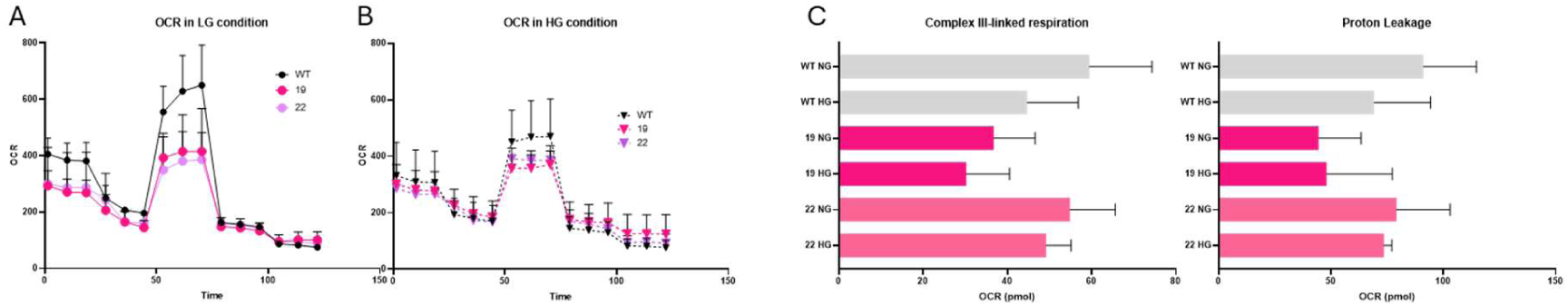
Metabolism assessment of *Ndr2* downregulated BV-2 cultured cells. Mitochondrial oxygen consumption of WT and *Ndr2* downregulated BV-2 exposed to (A) normal glucose conditions and (B) high glucose conditions. (C) Effect of Ndr2 downregulation on the complex III-linked respiration, measured after injection of rotenone and antimycin A, and on the proton leakage, measured after the injection of oligomycin and before the injection of BAM15, by the Seahorse assay (n=3-4).

**Supp. Figure S5.**
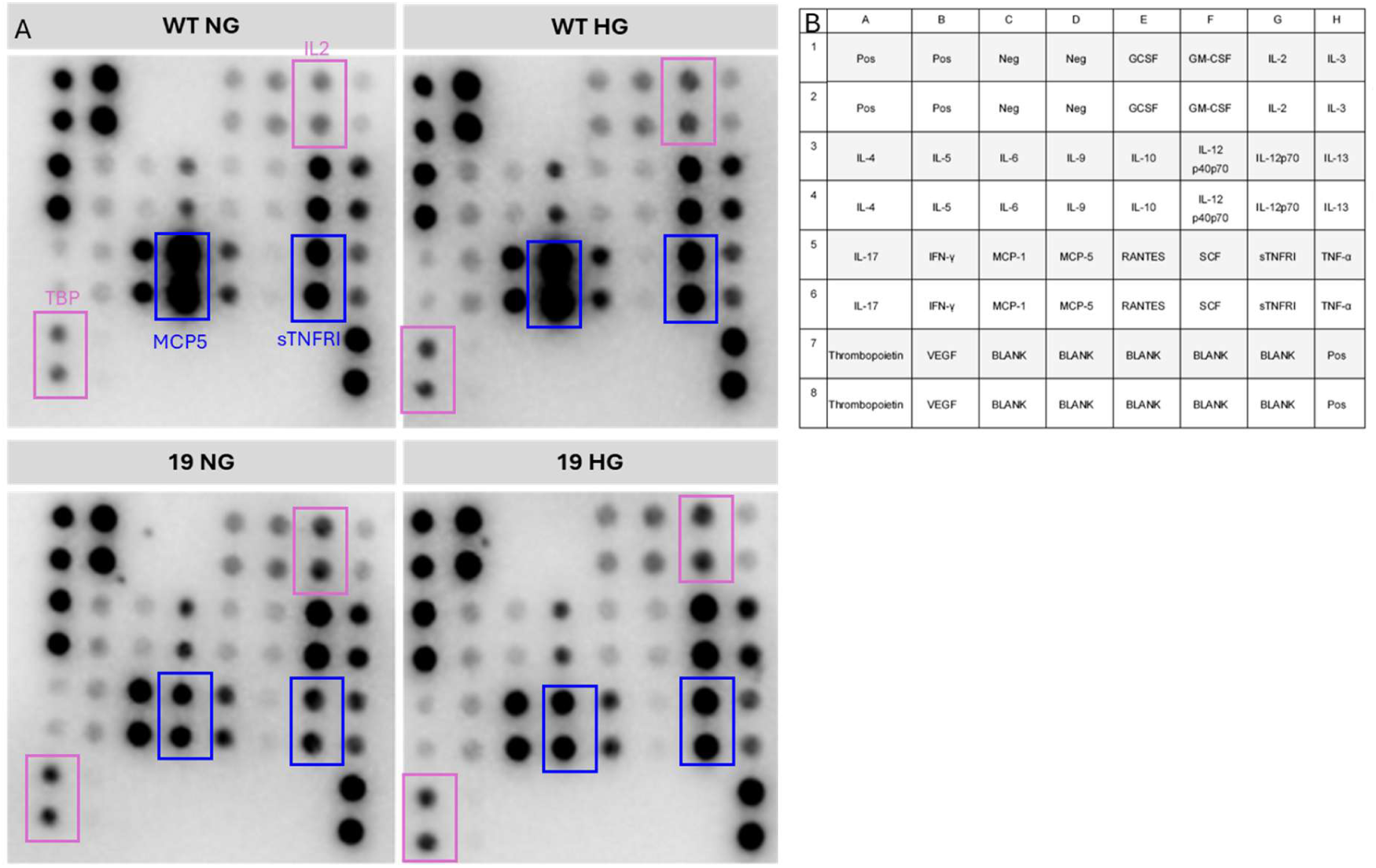
Expression of inflammatory factors in WT and *Ndr2* downregulated BV-2 under control and high glucose conditions. (A) Images of the cytokine array membranes exposed to the conditioned-media of WT normal glucose (WT NG or CT), WT high glucose (WT HG), and *Ndr2* downregulated clone 19 BV-2 cells, exposed to normal or high glucose (19 NG and 19 HG), respectively. (B) Map of the Mouse Cytokine Antibody Array (Membrane, 22 Targets), Abcam reference ab133993.

## References

1. Cabrera AP, Monickaraj F, Rangasamy S, Hobbs S, McGuire P, Das A (2020) Do genomic factors play a role in diabetic retinopathy? J Clin Med 9

2. Santiago AR, Boia R, Aires ID, Ambrósio AF, Fernandes R (2018) Sweet Stress: Coping with vascular dysfunction in diabetic retinopathy. Front Physiol

3. Rashid K, Akhtar-Schaefer I, Langmann T (2019) Microglia in retinal degeneration. Front Immunol 10(AUG):474669. 10.3389/FIMMU.2019.01975/BIBTEX

4. Altmann C, Schmidt MHH (2018) The role of microglia in diabetic retinopathy: Inflammation, microvasculature defects and neurodegeneration. Int J Mol Sci

5. Hergovich A (2016) The roles of NDR protein kinases in hippo signalling. Genes (Basel) 7(5):1–16. 10.3390/genes7050021

6. Hergovich A, Stegert MR, Schmitz D, Hemmings B a (2006) NDR kinases regulate essential cell processes from yeast to humans. Nat Rev Mol Cell Biol 7(4):253–64. 10.1038/nrm1891

7. Hergovich A (2013) Regulation and functions of mammalian LATS / NDR kinases : looking beyond canonical Hippo signalling. 1–12

8. Sharif AAD, Hergovich A (2018) Seminars in Cancer Biology The NDR / LATS protein kinases in immunology and cancer biology. Semin Cancer Biol 48(December 2016):104–114. 10.1016/j.semcancer.2017.04.010

9. Ardestani A, Lupse B, Maedler K (2018) Hippo Signaling: Key Emerging Pathway in Cellular and Whole-Body Metabolism. Trends in Endocrinology and Metabolism

10. Ye X, Ong N, An H, Zheng Y (2020) The Emerging Roles of NDR1/2 in Infection and Inflammation. Front Immunol 11:534. 10.3389/fimmu.2020.00534

11. Mia MM, Singh MK (2022) Emerging roles of the Hippo signaling pathway in modulating immune response and inflammation-driven tissue repair and remodeling. FEBS J 289(14):4061–4081. 10.1111/febs.16449

12. Léger H, Santana E, Leu NA, et al (2018) Ndr kinases regulate retinal interneuron proliferation and homeostasis. Sci Rep. 10.1038/s41598-018-30492-9

13. Goldstein O, Kukekova A V., Aguirre GD, Acland GM (2010) Exonic SINE insertion in STK38L causes canine early retinal degeneration (erd). Genomics 96(6):362–368. 10.1016/j.ygeno.2010.09.003

14. Berta ÁI, Boesze-Battaglia K, Genini S, et al (2011) Photoreceptor cell death, proliferation and formation of hybrid rod/S-cone photoreceptors in the degenerating STK38L mutant retina. PLoS One 6(9). 10.1371/journal.pone.0024074

15. Gardiner KL, Downs L, Berta-Antalics AI, Santana E, Aguirre GD, Genini S (2016) Photoreceptor proliferation and dysregulation of cell cycle genes in early onset inherited retinal degenerations. BMC Genomics. 10.1186/s12864-016-2477-9

16. Appelbaum T, Santana E, Aguirre GD (2017) Strong upregulation of inflammatory genes accompanies photoreceptor demise in canine models of retinal degeneration. PLoS One 12(5):e0177224–e0177224. 10.1371/journal.pone.0177224

17. Santos PF, Fazendeiro B, Luca FC, Ambrósio AF, Léger H (2023) The NDR/LATS protein kinases in neurobiology: Key regulators of cell proliferation, differentiation and migration in the ocular and central nervous system. Eur J Cell Biol 102(2):151333. 10.1016/J.EJCB.2023.151333

18. Zhang Z, Hou H, Yu S, et al (2020) Inflammation-induced mammalian target of rapamycin signaling is essential for retina regeneration. Glia 68(1):111–127. 10.1002/glia.23707

19. Lepiarz-Raba I, Gbadamosi I, Florea R, Paolicelli RC, Jawaid A (2023) Metabolic regulation of microglial phagocytosis: Implications for Alzheimer’s disease therapeutics. Transl Neurodegener 12(1):48. 10.1186/S40035-023-00382-W

20. Bernier L-P, York EM, Kamyabi A, Choi HB, Weilinger NL, MacVicar BA (2020) Microglial metabolic flexibility supports immune surveillance of the brain parenchyma. Nat Commun 11(1):1559. 10.1038/s41467-020-15267-z

21. LI L, QU C, WANG F (2015) A novel method for co-culture with Müller cells and microglia in rat retina in vitro. Biomed Rep 3(1):25–27. 10.3892/BR.2014.370

22. Aires ID, Boia R, Rodrigues-Neves AC, et al (2019) Blockade of microglial adenosine A2A receptor suppresses elevated pressure-induced inflammation, oxidative stress, and cell death in retinal cells. Glia 67(5):896–914. 10.1002/GLIA.23579

23. Haenseler W, Sansom SN, Buchrieser J, et al (2017) A Highly Efficient Human Pluripotent Stem Cell Microglia Model Displays a Neuronal-Co-culture-Specific Expression Profile and Inflammatory Response. Stem Cell Reports 8(6):1727. 10.1016/J.STEMCR.2017.05.017

24. Ran FA, Hsu PD, Wright J, Agarwala V, Scott DA, Zhang F (2013) Genome engineering using the CRISPR-Cas9 system. Nature Protocols 2013 8:11 8(11):2281–2308. 10.1038/nprot.2013.143

25. Labun K, Montague TG, Gagnon JA, Thyme SB, Valen E (2016) CHOPCHOP v2: a web tool for the next generation of CRISPR genome engineering. Nucleic Acids Res 44(W1):W272–W276. 10.1093/NAR/GKW398

26. Montague TG, Cruz JM, Gagnon JA, Church GM, Valen E (2014) CHOPCHOP: a CRISPR/Cas9 and TALEN web tool for genome editing. Nucleic Acids Res 42(Web Server issue). 10.1093/NAR/GKU410

27. Whale AS, De Spiegelaere W, Trypsteen W, et al (2020) The Digital MIQE Guidelines Update: Minimum Information for Publication of Quantitative Digital PCR Experiments for 2020. Clin Chem 66(8):1012–1029. 10.1093/CLINCHEM/HVAA125

28. Aires ID, Ribeiro-Rodrigues T, Boia R, et al (2020) Exosomes derived from microglia exposed to elevated pressure amplify the neuroinflammatory response in retinal cells. Glia 68(12):2705–2724. 10.1002/glia.23880

29. Jorge J, Neves J, Alves R, Geraldes C, Gonçalves AC, Sarmento-Ribeiro AB (2023) Parthenolide Induces ROS-Mediated Apoptosis in Lymphoid Malignancies. International Journal of Molecular Sciences 2023, Vol 24, Page 9167 24(11):9167. 10.3390/IJMS24119167

30. Schumacher TJ, Iyer A V., Rumbley J, Ronayne CT, Mereddy VR (2024) Exploring the impact of mitochondrial-targeting anthelmintic agents with GLUT1 inhibitor BAY-876 on breast cancer cell metabolism. BMC Cancer 24(1):1415. 10.1186/S12885-024-13186-6

31. Baptista FI, Aveleira CA, Castilho ÁF, Ambrósio AF (2017) Elevated Glucose and Interleukin-1β Differentially Affect Retinal Microglial Cell Proliferation. Mediators Inflamm 2017:4316316. 10.1155/2017/4316316

32. Yu F-X, Guan K-L (2013) The Hippo pathway: regulators and regulations. Genes Dev 27(4):355–371. 10.1101/gad.210773.112

33. Tamaskovic R, Bichsel SJ, Hemmings BA, Ã BAH (2003) NDR family of AGC kinases essential regulators of the cell cycle and morphogenesis. FEBS Lett 546(1):73–80. 10.1016/S0014-5793(03)00474-5

33. Tamaskovic R, Bichsel SJ, Hemmings BA, (2003) NDR family of AGC kinases essential regulators of the cell cycle and morphogenesis. FEBS Lett 546(1):73–80. 10.1016/S0014-5793(03)00474-5

34. Hsieh C-F, Liu C-K, Lee C-T, Yu L-E, Wang J-Y (2019) Acute glucose fluctuation impacts microglial activity, leading to inflammatory activation or self-degradation. Sci Rep 9(1):840. 10.1038/s41598-018-37215-0

35. Orihuela R, McPherson CA, Harry GJ (2016) Microglial M1/M2 polarization and metabolic states. Br J Pharmacol 173(4):649–665. 10.1111/bph.13139

36. Fairley LH, Lai KO, Wong JH, et al (2023) Mitochondrial control of microglial phagocytosis by the translocator protein and hexokinase 2 in Alzheimer’s disease. Proc Natl Acad Sci U S A 120(8). 10.1073/PNAS.2209177120

37. Okpala P, Omenyi S, Ozoegwu G, Achebe C (2015) Analysis of the dynamic energy flow associated with phagocytosis of bacteria. Heliyon 1(1):e00021. 10.1016/J.HELIYON.2015.E00021

38. Cheng J, Zhang R, Xu Z, et al (2021) Early glycolytic reprogramming controls microglial inflammatory activation. J Neuroinflammation 18(1):129. 10.1186/S12974-021-02187-Y

39. Li Y, Long W, Gao M, et al (2021) Trem2 regulates high glucose-induced microglial inflammation via the nlrp3 signaling pathway. Brain Sci 11(7):896. 10.3390/BRAINSCI11070896/S1

40. Kongtawelert P, Kaewmool C, Phitak T, Phimphilai M, Pothacharoen P, Shwe TH (2022) Sesamin protects against neurotoxicity via inhibition of microglial activation under high glucose circumstances through modulating p38 and JNK signaling pathways. Scientific Reports 2022 12:1 12(1):1–16. 10.1038/s41598-022-15411-3

41. Kinuthia UM, Wolf A, Langmann T (2020) Microglia and Inflammatory Responses in Diabetic Retinopathy. Front Immunol 11:564077. 10.3389/FIMMU.2020.564077/PDF

42. Hu A, Schmidt MHH, Heinig N (2024) Microglia in retinal angiogenesis and diabetic retinopathy. Angiogenesis 27(3):311. 10.1007/S10456-024-09911-1

